# Intergenerational effects of a paternal Western diet during adolescence on offspring gut microbiota, stress reactivity and social behavior

**DOI:** 10.1101/2021.09.16.460599

**Authors:** Carina Bodden, Terence Y. Pang, Yingshi Feng, Faria Mridha, Geraldine Kong, Shanshan Li, Matthew J. Watt, Amy C. Reichelt, Anthony J. Hannan

## Abstract

The global consumption of highly processed, calorie-dense foods has contributed to an epidemic of overweight and obesity, along with negative consequences for metabolic dysfunction and disease susceptibility. As it becomes apparent that overweight and obesity have ripple effects through generations, understanding of the processes involved is required, in both maternal and paternal epigenetic inheritance. We focused on the patrilineal effects of a Western-style high-fat (21%) and high-sugar (34%) diet (WD) compared to control diet (CD) during adolescence and investigated F0 and F1 mice for physiological and behavioral changes. F0 males (fathers) showed increased body weight, impaired glycemic control, and decreased attractiveness to females. Paternal WD caused significant phenotypic changes in F1 offspring, including higher body weights of pups, increased Actinobacteria abundance in the gut microbiota (ascertained using 16S microbiome profiling), a food preference for WD pellets, increased male dominance and attractiveness to females, as well as decreased behavioral despair. These results collectively demonstrate the long-term intergenerational effects of a Western-style diet during paternal adolescence. The behavioral and physiological alterations in F1 offspring provide evidence of adaptive paternal programming via epigenetic inheritance. These findings have important implications for understanding paternally mediated intergenerational inheritance, and its relevance to offspring health and disease susceptibility.

## Introduction

In evolutionary terms, the overconsumption of energy-dense foods as well as a sedentary lifestyle are very recent problems of our modern society ^1, 2^. Epidemiological studies have shown that ultra-processed, calorie-dense foods promote the development of overweight and obesity – defined as abnormal or excessive fat accumulation that may impair health ^1, 3^. Alarmingly, the prevalence rate of obesity in adults increased substantially over the past three decades and has reached epidemic proportions ^4^. Hence, the healthcare system and society faces an enormous clinical and economic burden as overweight and obesity promote negative health consequences, such as cardiovascular disease, type 2 diabetes, several cancers, and a diminished life expectancy ^5^.

Although the adverse effects of a Western-style diet on the body, including brain function have been studied closely in recent years, there are numerous remaining questions. One reason is the fact that human studies are confounded by additional factors, such as genetic predisposition, exercise ^6^ and smoking ^7^, rendering the relationship between diet and health effects more complicated. To understand how specific nutritional factors can impact on an individual’s physical and mental health, other parameters need to be controlled. Animal models provide the opportunity to rigorously control for genetic as well as environmental factors, including nutritional factors. The shorter lifespan of mice, for example, also brings about numerous advantages compared to long-lived organisms. Long-term effects can be observed sooner and nutritional effects during particular life stages can be studied more easily. The short generation time furthermore renders the possibility to look at intergenerational and transgenerational effects.

Since there is evidence for generation-spanning effects of Western diets, rodent studies provide an excellent basis for the study of these effects ^8^. Inter- and transgenerational inheritance are mediated via epigenetic modifications of the germ cells. This phenomenon, encapsulated in the Developmental Origins of Health and Disease (DOHaD) research framework, assumes that early-life environmental factors, such as parental diet, can increase the susceptibly of the offspring to diseases, such as metabolic disturbances, later in life ^9^. Maternal ‘programming’ has already been studied intensively, whereas the influence of paternal programming has received relatively little attention. Yet, recent evidence demonstrates that diet-induced alterations in epigenetic markers in sperm that can influence the offspring phenotype ^8, 10^. It remains unclear the extent to which various aspects of offspring phenotypes can be modified by diet, and whether sperm modifications are acutely or permanently influenced by diet.

Males start to produce spermatozoa around the onset of adolescence. Children and adolescents are particularly prone to consume unhealthy diets, which has contributed to a tenfold increase of obesity in children over the past four decades ^1^. That the developing brain is plastic and can be influenced by nutritional factors is well acknowledged ^11^ and recent work shows that one week of an acute high-sugar intake is sufficient to negatively affect sperm quality ^12^, indicating a high susceptibility to environmental influences. Therefore, the question arises as to whether an unhealthy diet around the onset of spermatogenesis can lead to even more deleterious, potentially permanent effects? In this regard, most research has focused on the consequences of obesity, however, our understanding of the effects of unhealthy eating can have, prior to induction of obesity or diabetes, is limited. In rats, feeding a high-fat diet during the peri-pubertal period was linked to long-term effects on metabolism and the reproductive system ^13^. Specifically, high-fat diet fed rats displayed glucose intolerance, increased fat tissue deposition, reduced energy expenditure, an increase in the number of abnormal seminiferous tubules and a reduction in seminiferous epithelium height and seminiferous tubular diameter, and a decreased number of spermatozoa ^13^. The finding that diet can influence testis physiology and sperm count suggests an increased sensitivity of the reproductive organs including germ cells to environmental factors. This sensitivity could offer an evolutionary advantage by providing a mechanism whereby information about the current environmental exposures can be transferred to the offspring through paternal programming. More precisely, in the event of food abundance or food scarcity, the next generation could greatly benefit from potential adaptive processes that might provide a survival advantage in the prevailing environment.

Understanding the consequences of an unhealthy diet during sensitive phases of life, including the intergenerational impacts, is of utmost importance for the comprehensive understanding and prevention of nutrition-related diseases. Hence, the main aim of this study was to elucidate the intergenerational effects of a paternal Western-style diet during adolescence on physiology and behavior of F1 males and females. For a comprehensive analysis, we additionally investigated the influence of an adolescent Western diet on physiology and behavior of F0 males.

## Materials and Methods

### Aims

The main aim of this study was to elucidate the intergenerational effects of a paternal Western-style diet during adolescence on physiology and behavior of F1 males and females. For a comprehensive analysis, we additionally investigated the influence of an adolescent Western diet on physiology and behavior of F0 males.

### Animals and housing conditions

C57BL/6J breeder mice were purchased from the Animal Resources Centre (Murdoch, WA, Australia) and arrived at the Florey Institute at 4 weeks of age, when they were randomly assigned to one of two experimental groups: Western-style diet (WD; #SF00-219, Specialty Feeds, Glen Forrest, WA, Australia) or control diet (CD; #AIN93G, Specialty Feeds, Glen Forrest, WA, Australia). WD or CD exposure started on the day of arrival (**Figure 1**). Male mice were housed in groups of 2-5 animals until postnatal day (PND) 70, when they were pair-mated with nulliparous female C57BL/6J mice for 5 days. The females were previously group-housed in same-sex cages and fed standard chow (#102108, Barastoc, Ridley, VIC, Australia). Afterwards, males were individually housed and reintroduced to their respective diet. Another subgroup of F0 males was not mated but instead tested for differences in glucose tolerance and their behavioral and physiological phenotype at 10 weeks of age. F1 Offspring were weaned at 4 weeks of age and housed in same-sex groups of 2 to 5 animals from the same paternal treatment (paternal CD, PatCD or paternal WD, PatWD).

**Figure 1.**
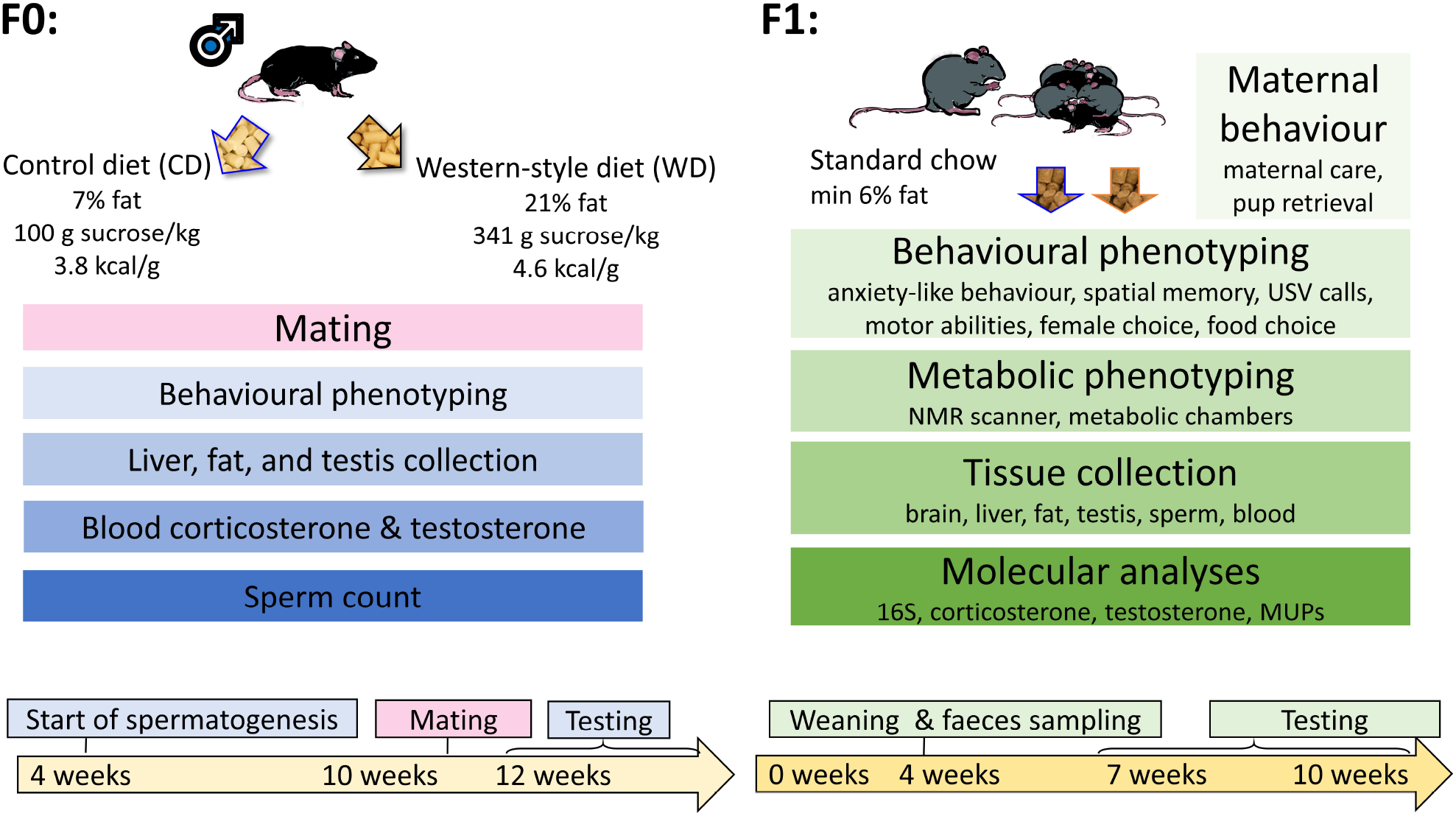
Experimental design. Male C57BL/6J mice were fed either a control (CD) or a Western-style diet (WD) from 4 weeks of age onwards. These F0 males were mated with standard-fed female C57BL/6J mice. F0 males and F1 offspring were assessed for behavioral, physiological, and endocrinological changes.

All animals were group-housed (up to five per cage) in standard open-top mouse cages (15 cm × 30 cm × 12 cm) with food and filtered tap water *ad libitum*. Cages were equipped with wood chip bedding and two tissues as nesting materials and were cleaned on a weekly basis. Mice were housed in the Florey Institute animal facility in a room maintained at a temperature of 22-25°C and 55% humidity on a 12:12 h light–dark cycle, with lights on at 7 am).

### Experimental design

#### Paternal diet

Diet manipulation of male F0 mice commenced at 4 weeks of age (postnatal day 28) and continued for 6-10 weeks (depending on experimental subgroup), which encompassed adolescence and puberty ^14^. According to the latest recommendations ^15^, commercial WD and CD chows were acquired and used to ensure comparability with other studies (e.g., ^16, 17^). WD (#SF00-219, Specialty Feeds, Glen Forrest, WA, Australia; Harlan Teklad TD 88137 equivalent, intended to mimic a Western-style fast-food diet) comprised of 21% by weight saturated fat (clarified ghee) and 34% sucrose (digestible energy: 19.4 MJ/Kg). The CD (#AIN93G, Specialty Feeds, Glen Forrest, WA, Australia) contained 7% fat and 10% sugar (digestible energy: 16.1 MJ/Kg. Protein (19%) and vitamin contents were comparable in both diets.

#### Breeding

At 10 weeks of age, male F0 mice were pair-mated with nulliparous females for 5 days. During the mating period, all mice were fed standard chow. Afterwards, female mice were separated to litter down undisturbed in single-housing cages. Non-pregnant females and females who lost their litters were excluded from the study. Litter size was not adjusted. At 4 weeks of age, F1 offspring were weaned and housed in same-sex groups of 2-5 animals from the same paternal treatment (PatCD or PatWD).

#### Maternal behavior

Maternal care in the home cage was observed daily starting at birth (PND 0) until PND 10. All behavior observations were performed by an experienced observer (C.B.) using instantaneous sampling, which refers to the recording of an individual’s behavior at sequential, predetermined points in time (20 s intervals). Two observation sessions were conducted per day, one between 10:00 am and 11:00 am and one between 3:00 pm and 4:00 pm. 30 scans were performed for each animal in one observation session; therefore 60 scans were conducted per day per animal. The order of animals was randomized but was the same for each observation session. For data analysis, the percentage of intervals, in which each behavior occurred was calculated. (Descriptions of maternal behaviors are summarized in **Suppl. Table 1**.)

#### Pup retrieval

One randomly selected pup was carefully removed from the nest in its home cage and placed in the corner opposite of the nest. The time until the dam reached the pup and picked it up was considered the latency to retrieve the pup. This procedure was repeated five times with different pups and the mean time to retrieve pups was calculated.

### Physiological parameters

#### Body weight

Body weights of male F0 breeders and F1 offspring were monitored weekly using a digital scale.

#### Glucose tolerance test

A glucose tolerance test (GTT) was performed in F0 males at the age of 10 weeks. Mice were fasted for 14 hours overnight. Body weight and fasting glucose levels were determined by placing each mouse into a restrainer, puncturing a tail vein and collecting blood directly into the glucometer (Accu-chek, Roche). Subsequently, a sterile glucose solution (10% (w/v) in PBS) was administered by intraperitoneal injection at a concentration of 1 g/kg (glucose/body weight). Blood samples were taken at 15, 30, 60, and 90 min thereafter. In between blood collections, mice were released into their home cages.

#### Metabolic testing

For metabolic testing, a subgroup of F1 animals (N = 6 – 10) was transferred to the Melbourne Mouse Metabolic Phenotyping Platform (MMMPP, Department of Physiology, University of Melbourne) and allowed to habituate for at least 7 days prior to testing.

#### Metabolic cages

The Promethion Metabolic System (Sable, Nevada, USA) was used to measure food intake, water intake, activity, oxygen consumption and carbon dioxide production in individual mice over 48 h, following an acclimatization period of 12 h. The metabolic cages were approximately the same size as the normal home cages and were equipped with food, water, and bedding. Subsequently, mice were returned to their original home cages for 7 days prior to tissue collection/endpoint.

#### Body composition

Body composition (fat mass, lean mass, and free fluid) was measured by the Bruker LF50 MiniSpec (Billerica, Massachusetts, USA), which uses Time Domain Nuclear Magnetic Resonance (TD-NMR) in conscious mice without the need for anesthesia. Scans were completed in approximately 2 min after which mice were released to their home cage.

#### Blood, tissue, and sperm collection

##### Plasma collection

Mice were deeply anesthetized using isoflurane (4-5% in O_2_) for blood collection through cardiac puncture, immediately followed by cervical dislocation. Blood was collected in EDTA-coated tubes (Sarstedt) before separation of plasma by centrifugation at 1100 g for 10 min. Plasma was then stored at −20°C until subsequent analyses.

##### Tissue collection

Testes, liver, posterior subcutaneous and epididymal fat were removed and weighed.

##### Sperm collection

As described previously ^18^, cauda epididymides were separated from the corpus segment and the epididymal fat pad was removed, leaving only the vas deferens attached. The cauda epididymides were bisected with a single incision, then immersed into 1 ml of pre-warmed (37°C) M2 medium (M7167, Sigma-Aldrich, NSW, Australia). After an incubation phase of at least 30 min at 37°C, tissue was discarded, and spermatozoa were resuspended. An aliquot was removed and stained with Trypan Blue for the purpose of cell counting.

#### Histological analysis

##### Testes, liver, and posterior subcutaneous fat

Tissues were formalin-fixed (10%), dehydrated, paraffin embedded, sectioned at 4 μm, and stained with hematoxylin and eosin (H&E). Six randomly selected sections per mouse were examined under a light microscope (#BX61, Olympus, Tokyo, Japan) and photomicrographs were captured with Image-Pro Plus 6.0 (Media Cybernetics, Rockville, MD, USA). The histological analysis was performed using ImageJ software (National Institutes of Health, Bethesda, MD, USA). Lipid droplet area and adipocyte size were quantified using the automatic particle counting plugin of ImageJ to avoid a user bias. Manual counts were performed to verify the accuracy of the automatic analysis. Testis parameters (**see Figure 4I**) were manually assessed by an experimenter blind to experimental groups. The volumetric density of testes was determined according to the protocol published previously ^19^.

#### Testosterone and corticosterone

##### Testosterone analysis

Plasma samples were diluted to 1:30 and assayed using a testosterone ELISA according to the manufacturer’s instructions (#582701, Cayman Chemical, Ann Arbor, MI, USA). For measurements of testicular testosterone levels, one testis per mouse was lysed in 600 μl 10 mM PBS, sonicated (4°C for 2 × 20 s) and centrifuged (10,000 g at 4°C for 10 min). A small aliquot of the supernatant (10 μl) was then purified from 2 ml diethyl ether under a gentle stream of nitrogen gas. Pellets were reconstituted with 50 μl assay buffer and further diluted to 1:25 and 1:100 for the testosterone ELISA.

##### Corticosterone analysis

Plasma samples were diluted 1:10 for control (unstressed) mice and 1:500 for stressed mice (15 min after Porsolt swim test). Corticosterone was assessed using an EIA as per manufacturer’s instructions (#501320, Cayman Chemical, Ann Arbor, MI, USA).

#### MUPS quantification

##### Urine sampling

Urine was collected in 1.5 ml tubes directly from the mice as they were tail and scruff handled. Samples were immediately frozen on dry ice and stored at –80°C.

##### Protein quantification

Protein concentration of urine samples was measured according to a standard Bradford assay on a 96-well microplate. Total (MUP concentration, see below) and relative (protein/creatinine ratio) urinary protein concentration was evaluated, corrected for urine dilution by measuring creatinine (Creatinine urinary colorimetric assay kit, Cayman Chemical, Ann Arbor, MI, USA). Since the vast majority of urinary proteins are MUPs (up to 99%; ^20, 21^), total urinary protein concentration was used to estimate MUP concentration.

### Gut microbiome analysis

#### Fecal collection

When F1 offspring were 28 postnatal days of age (4 weeks old), each was put into a new clean cage (wiped with 80% ethanol) until at least two fecal pellets were excreted. Fresh pellets were collected, frozen immediately on dry ice (–80°C) and transported to the Peter Doherty Institute for Infection and Immunity (University of Melbourne, Melbourne, Australia) on dry ice for microbiome analysis.

#### Microbiome analysis

Briefly, genomic DNA of fecal pellets was extracted using the Qiagen Powersoil Kit (Qiagen, Germantown, MD, USA). The bacterial V4 16S ribosomal RNA gene was amplified using 515 forward barcoded and 806 reverse primers following the Earth Microbiome Project ^22^. Next, amplicons were sequenced using Illumina MiSeq (Illumina, Inc., San Diego, CA, USA). Raw FASTQ sequences were processed using Qiita, including sequence quality control, demultiplexing, trimming and resolving amplicon sequence variants (ASVs) with Deblur ^23^. The representative sequences were mapped onto the Greengenes reference database (gg-13-8-99-515-806-nb-classifier) to obtain the taxonomic identity of ASVs. The count table consisting of 759 ASVs were further analyzed for their biodiversity using QIIME2-2019.7 ^24^. For the subsequent analysis, the data was first stratified by sex to examine sex-specific effects on microbiome. As there were no differences between the sexes, they were combined to examine the overall effect of paternal diet on the microbiome.

### Behavioral testing

If not stated differently, behavioral tests were performed between the age of 7 to 12 weeks of age, with a recovery time of at least 24h between each test. Behavioral testing was conducted in a room separate from the housing room. All mice were allowed to acclimatize to the testing room for at least 30 min prior to commencement of each test. Testing was conducted between 0800 and 1600 h under dimmed lighting of 25–50 lx. Animals were tested in a randomized order with the experimenter blind to the paternal condition of F1 mice. The sequence of tests was the same for all mice. Testing arenas were thoroughly cleaned with 80% ethanol between tests. An overhead video camera was used to record the animals’ movements. Behaviors were analyzed using the CleverSys TopScan tracking software (Reston, VA, USA). (Behavioral tests conducted in F0 and F1 generations are summarized in **Suppl. Table 2**.)

#### Neonatal testing

##### Surface righting

On PND 7, F1 pups were tested for motor function, where they were placed on their back on an even surface, held in that position for 5 s and then the time they needed to independently turn over and stand upright on their feet was assessed (max. 60 s).

##### Ultrasonic vocalization (USV) recording

On PND 8, F1 pups were carefully removed from the home cage and individually placed into a small open top container for 2 min. An ultrasound Microphone (Avisoft UltraSoundGate condenser microphone capsule CM16, Avisoft Bioacoustics, Berlin, Germany) sensitive to frequencies between 10 – 180 kHz was situated approximately 20 cm above the center of the container to measure any USVs emitted by the pup during testing. USVs were subsequently recorded using Avisoft Recorder software (Version 3.2). Settings included a sampling rate of 250 kHz and a resolution of 16 bits. For the analysis, recordings were transferred to Avisoft SASLab Pro (Version 5.2.07) and a fast Fourier transformation (FFT) was conducted. Spectrograms were generated with an FFT-length of 512 points and a temporal window overlap of 50% (100% Frame, FlatTop window). The spectrogram was produced at a frequency resolution of 488 Hz and a temporal resolution of 1 msec. A lower cut-off frequency of 20 kHz was used to remove all audible background noise. Spectrograms were visualized up to 125 kHz. The latency to call and the call frequency were assessed.

#### Anxiety-like and exploratory behavior

##### Elevated-plus maze

The test was performed to assess anxiety-like and exploratory behavior as previously described ^25^. Briefly, the apparatus consisted of a plus-shaped maze, with 4 arms extending from the center (5 cm × 5 cm). Arms were 30 cm in length and 5 cm in width and the closed arms had surrounding walls of 14 cm height. Mice were allowed to freely explore the maze for 5 min. The time spent on the open arms was expressed as percentage of the sum of the time on open and closed arms. Furthermore, unprotected and protected head dips were assessed manually from video recordings as a measure of risk assessment.

##### Light-dark box

The test was used as another assessment of anxiety-like and exploratory behavior and performed as previously described ^25^. Briefly, the test was conducted using the Med Associates ENV-510/511 open-field arenas (Fairfax, VT, USA). The light compartment was illuminated with overhead LED lamps to 750 lx. Mice started the 10 min test session in the dark insert. Time spent in each half of the arena was automatically tabulated by the proprietary activity monitor software and expressed as a percentage of the test duration.

#### Learning and memory

##### Novel object recognition (NOR)

The test was performed to assess object memory and was adapted from Leger and colleagues ^26^. Mice were habituated to the empty testing arena (50 cm x 50 cm) for 10 min. On the next day, the mice were individually placed in the same testing arena, in which two novel but identical objects (transparent plastic flasks filled with sand or colorful plastic Lego towers of approximately same size but different shape and color) and allowed to freely explore for 10 min. Both objects and location were counterbalanced to avoid any effect of object and/or location preference. After an inter-trial interval of 1 h, animals were placed back into the testing area that now contained one novel object as well as one familiar object. Animals were allowed to explore the arena for 10 min. The arena and objects were cleaned with 10% ethanol between trials to remove odor cues.

##### Y-maze

The apparatus consisted of symmetrical arms (each 30 cm in length) with 14 cm high walls. The mouse was placed at the end of the ‘home arm’ and allowed to explore 2 of the 3 arms for 15 min. A partition blocking off the novel arm of the maze was in place during this initial trial. Each of the three arms was marked by a unique cue for spatial orientation. The second test trial was performed after an inter-trial interval of 1 h. During the 5 min test trial the partition was removed so that all arms were available for exploration.

#### Social interest, social recognition, and mate preference

##### Social interest and social recognition test

Mice were habituated to the three-chamber-arena containing two empty mesh wire cages for 10 min. In the following social interaction trial, one unfamiliar, same-sex, same-age animal was placed under one of the cages. The mouse was reintroduced to the testing arena and allowed to move freely and interact with the ‘stimulus mouse’ for 10 min. In the subsequent social recognition trial, a second unfamiliar same-sex, same-age mouse was placed under the second cage and the test subject was again allowed to explore to arena for 10 min.

##### Female preference

Female mice were assessed for the stage of their estrous cycle in the morning. Briefly, a small droplet of saline (around 20 µL) was gently inserted into the vagina with a P20 pipette and the lavage liquid was mounted on a slide for observation on a microscope under 10X magnification. The different cell type populations were analyzed to determine the estrous cycle stage (Byers et al., 2012). Females in estrous were transferred to a behavioral testing room for acclimatization of at least 30 min before testing commenced. The choice apparatus consisted of a three-chamber test with a middle chamber with access to two lateral chambers, both comprising a wire-mesh cage. In the habituation trial, the female was allowed to explore the three chambers with empty wire cages for 5 min. Afterwards, the female was removed, and the stimuli were placed inside the wire cages. These stimuli consisted of either a) male mice from different experimental groups (PatCD and PatWD) or b) a small amount of soiled bedding from the males’ original cage (bedding was considered soiled after at least 3 days after cleaning cages ^27^) or c) both male mice and bedding. The female was then re-introduced to the apparatus and allowed to explore the apparatus for 10 min. The percentage of time spent in each lateral chamber (preference) is considered as a measure of attractiveness ^28^. To avoid a side bias, the position of the wire cages and experimental groups was randomized.

#### Depression-like behavior

##### Saccharin preference

Mice were habituated to two water bottles filled with filtered tap water in their home cages one day before the test. During the test, mice had access to two bottles, one containing water and the other containing a saccharin solution (0.01%, diluted in filtered tap water). The position of the bottles was randomized to avoid a side bias. Fluid consumption was assessed by weighing both tubes at the beginning of the test and after 24 h. Saccharin intake and water intake were recorded for each cage. A saccharin preference index was calculated as saccharin solution intake/(saccharin solution intake + water intake).

##### Porsolt swim test

Mice were placed into a Perspex beaker (2.5 l) – filled to 1.7 l mark with water (23-25 °C). After 5 min, mice were removed from the water tank and placed back into their home cages. Videos were captured from the side and were used to assess the time that the animals spent swimming versus being immobile (floating) as a measure of depressive-like behavior and/or behavioral despair.

#### Food-motivated behavior and olfactory abilities

##### Novelty-suppressed feeding (NSF)

Before testing, mice were deprived of food for 24 h. The test was conducted under dimmed lighting of 25 lx in a novel arena (30 cm × 30 cm × 30 cm) with both a CD and a WD food pellet in the center. The latency for the mouse to assume a stationary posture while feeding on either pellet was recorded (max. 10 min) as was the choice of the pellet. When this occurred, the mouse was removed and returned to their original caging conditions.

##### Olfactory discrimination

This test was conducted to determine the ability of mice to discriminate between and habituate to different odor types. Mice were habituated to a room devoid of strong smells for a minimum of 30 min prior to testing. Testing occurred separately for each mouse in a standard cage with fresh bedding and a metal lid. The odors were presented on top of the metal lid in sequential order: water, vanilla, almond (cooking essence). Each odor was presented for a period of 2 min, repeated over 3 trials, with an inter-trial interval of 1 min. All odors were diluted at 1:4 in distilled water. At the beginning of every trial, 10 μL of the odor were pipetted onto a fresh piece of paper. The mouse stayed in the cage for the whole sequence of testing. Recorded video files were manually assessed for the time the mouse spent sniffing the odor source during each odor presentation by an observer blinded to the experimental groups. Sniffing was scored when the animal was orienting towards the odor source with its nose within 2 cm distance.

##### Food taste preference

Mice were provided with both WD and CD pellets in the food rack of their home cage for 48 h. Pellets were weighed before and after the test to assess a food preference. The higher porosity of WD pellets was associated with some food being dispersed in the bedding, requiring a thorough collection procedure.

##### Food smell preference

Two tea strainers were secured on either side of the metal lid of a standard cage. One strainer contained CD pellets, the other WD pellets. Sides were randomized between trials. Mice were individually placed into the cages and allowed to freely explore the strainers for 5 min. Using video files, the experimenter manually assessed how long the mice sniffed at each side. The experimenter was blinded to the type of diet and the experimental groups. Sniffing was defined as the nose being within the radius of 2 cm around the tea strainer.

##### Buried food test

Animals were habituated to cereals (Crunch®, Nestlé) on three consecutive days before the actual test. On test day, animals were fasted for 5 h. Testing occurred in 2 consecutive trials separated by a 3-min rest time. During the first trial, a piece of cereal was placed on the surface bedding at one randomly chosen location (out of 8), and the mouse was introduced in the center of the cage. The time to reach and start eating the cereal was measured with a manual timer and used to assess mice motivation to reach a food item. Mice were prevented from eating the entire piece of cereal to avoid satiation. The second trial was performed the same way, except that the cereal was buried beneath 2 cm of bedding. The test ended once either the animal found the cereal or did not find it within 10 min.

### Statistics

General linear models were used for data analysis. To meet the assumptions of parametric analysis, residuals were graphically examined for normal distribution, homoscedasticity and outliers, and the Shapiro-Wilk test was applied. When necessary, raw data were transformed using square root, logarithmic, or inverse transformations. Specifically, linear models were used to analyze dependent variables with fixed between-subject factors ‘paternal diet’ and ‘sex’. Main effects and interaction terms were tested on local significance level alpha (α) = 0.05. *Post-hoc* t-tests were applied, and Bonferroni-Holm corrected. F0 data was analyzed by t-tests. Since not all data were normally distributed and could not be transformed adequately, non-parametric statistics were applied. Specifically, the Mann-Whitney U test for between-group comparisons with Bonferroni-Holm correction was performed.

A statistical power analysis for sample size estimation was performed using G*Power ^29^. Taking into account all parameters that yielded large effect sizes, we could ensure that the total sample size could detect biologically relevant differences with a power of 80% ^30^. Statistical analyses were conducted using the statistical software IBM SPSS Statistics (IBM Version 26, Release 26.0.0.0). Graphs were created using the software GraphPad Prism (Version 8.3.0 (538) for Windows, GraphPad Software, San Diego, California USA).

## Results

### F0 generation

F0 male breeders were tested for differences in weight gain, anxiety-like behavior and locomotor activity, spatial learning, anhedonia, and female preference.

#### Physiologic parameters

There was a significant main effect of diet (F_(1,48)_ = 67.117, p < 0.001) and time (F_(2.2,103.5)_ = 886.423, p < 0.001) on body weight (**Figure 2A**). Furthermore, a significant interaction between diet and time (F_(2.2,103.6)_ = 36.351, p < 0.001) was detected. Body weight was significantly higher in males fed WD compared with CD. Furthermore, WD mice gained more weight over time than CD mice. Food consumption was significantly higher in WD compared to CD mice when assessed total mass (t = -6.756, p < 0.001; **Figure 2B**) or total energy intake (t = -11.490, p < 0.001; **Figure 2C**). In the GTT, a significant main effect of diet (F_(1,8)_ = 132.8, p < 0.001) and time (F_(1.873,14.98)_ = 36.54, p < 0.001) as well as a diet × time interaction (F_(4,32)_ = 7.349, p < 0.001) was detected, with mice fed the WD having an increased fasting blood glucose compared with mice fed the CD (t0: t = -8.816, p < 0.001; t15: t = -11.578, p < 0.001; t30: t = -4.535, p < 0.002, t60: t = -3.789, p = 0.005; **Figure 2D)**. However, there was no evidence for impaired glucose tolerance. WD males had a significantly higher epididymal fat mass (t = -3.109, p = 0.016; **Figure 2E**), higher posterior subcutaneous fat mass (t = -10.036, p < 0.001**; Figure 2F**) and greater adipocyte cell area (t = 4.983, p < 0.001; **Figure 2G, 2H**) compared to CD males. Liver weight was significantly increased in WD compared to CD (t = -4.216, p < 0.001; F**igure 2I**) and there was a clear evidence of hepatic steatosis in the WD group as indicated by a markedly increased total size of hepatic lipid droplets (t = 2.797, p = 0.042; **Figure 2J, 2K**) as well as percent coverage of sections (t = 2.467, p = 0.039).

**Figure 2.**
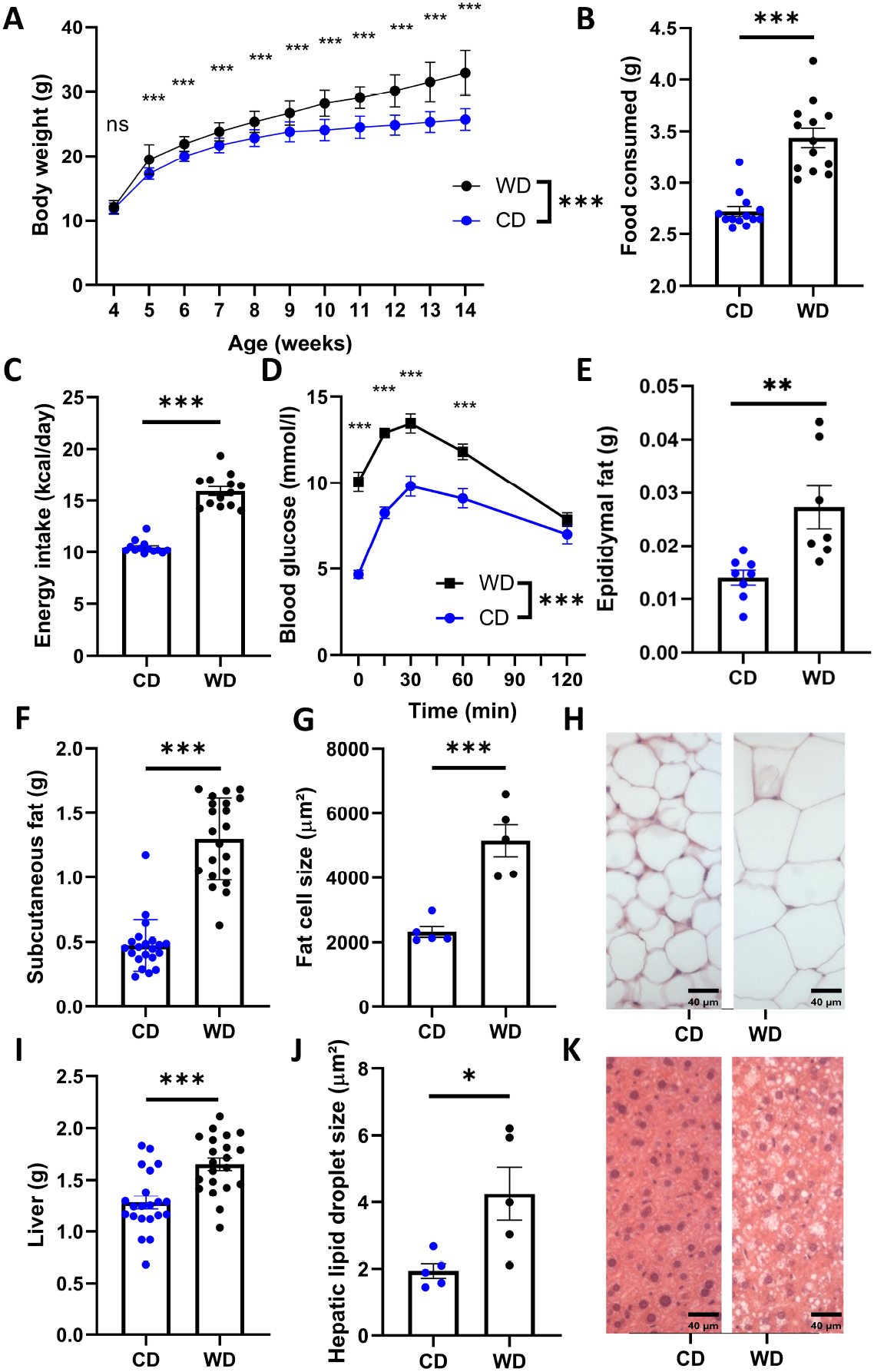
Physiological parameters of F0 males. **A)** Body weight development of F0 males fed either WD or CD (N = 25) as well as **B**) daily food consumption (N = 13), **C**) daily energy intake (N = 13), **D**) blood glucose level (N = 5)s, **E**) epididymal fat mass (N = 7-8), **F**) posterior subcutaneous fat mass (N = 21), **G**) adipocyte cell size (N = 5), **H**) adipocytes (H&E stained), **I**) liver mass (N = 21), **J**) hepatic lipid droplet size (N = 5), **K**) hepatic lipid droplets (H&E stained). Graphs show individual values where possible and means ± SEM. Statistics: General linear models (A, D), t-test (B-C, E-F, I-J). *, p < 0.05; **, p < 0.01; ***, p < 0.001.

#### Behavioral tests

Saccharin preference was significantly lower in WD compared to CD mice (t = 3.58, p = 0.002; **Suppl. Fig. 1**). In the mate-preference test, females did not show a preference when males were presented without soiled bedding in the wire-mesh cages (**Figure 3A**). However, when males were presented along with soiled bedding, females preferred the chamber of the CD males over the chamber of the WD males (t = 3.019, p = 0.005; **Figure 3B**). When only soiled bedding was provided in the cages, females still preferred the side of CD males over WD males (t = 4.907, p < 0.001; **Figure 3C**). CD compared to WD males showed more successful mountings during the mating session (U = 11.000, p = 0.003; **Figure 3D**).

**Figure 3.**
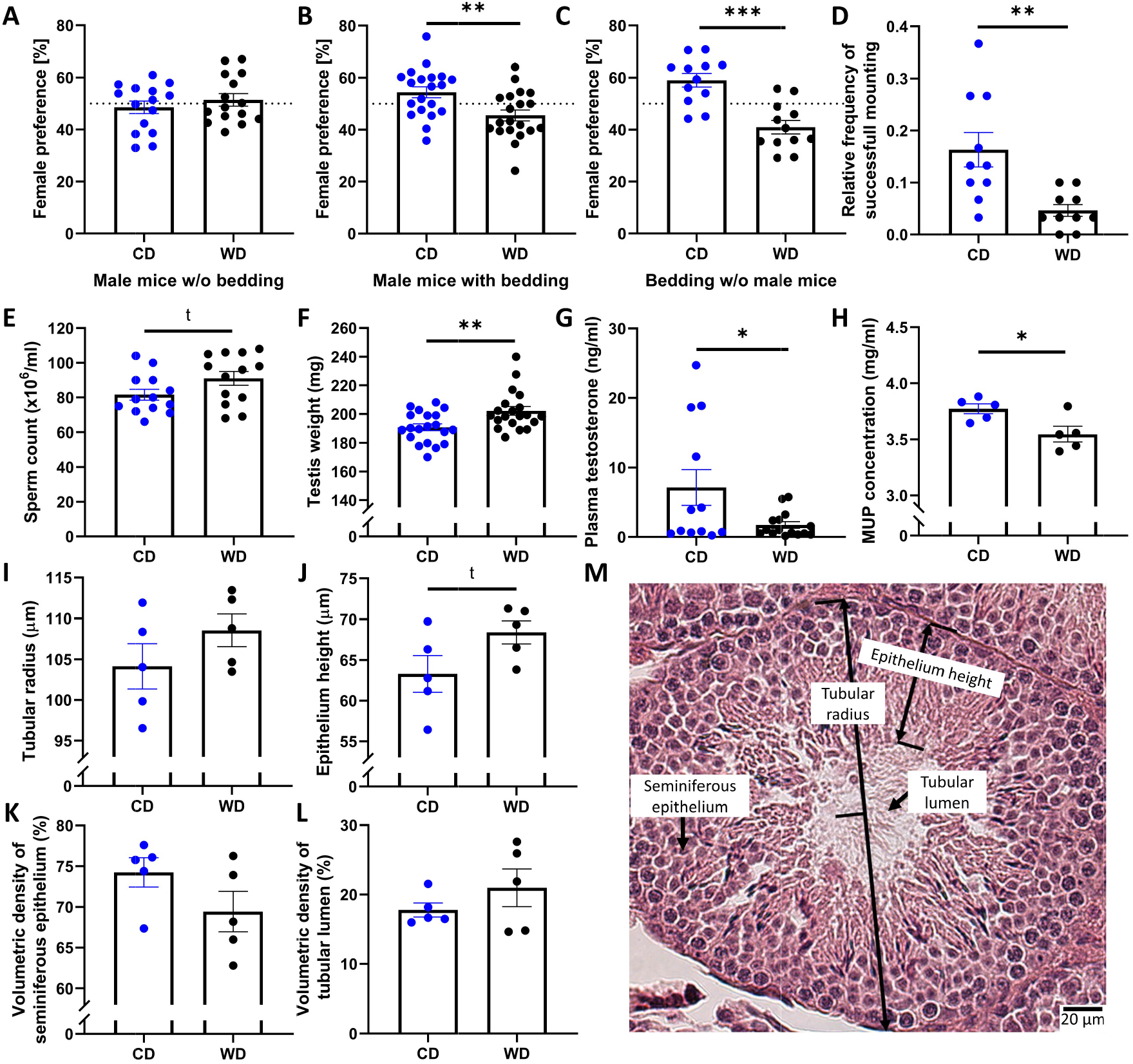
Attractiveness of F0 males and reproductive tract parameters. Results of female preference tests **A)** with CD and WD males without soiled bedding (N = 15), **B)** with CD and WD males with soiled bedding (N = 20), **C)** without CD and WD males but with soiled bedding (N = 12). **D)** Frequency of successful mounting in first hour of co-housing F0 males fed CD or WD with nulliparous females (N = 10). **E)** Sperm count (N = 13), **F)** Total testis weight (N = 20), **G)** plasma testosterone levels (N = 8), H) MUP concentration (N = 5), **I)** Tubular radius (N = 5) and **J)** epithelium height (N = 5). Volumetric density of **K)** seminiferous epithelium (N = 5) and **L)** tubular lumen (N = 5). **M)** depicts an example of an H&E stained testicular tubule including definitions of terms used. Graphs show individual values and means ± SEM. Statistics: t-test. *, p < 0.05; **, p < 0.01; ***, p < 0.001. Dotted lines indicate chance level.

There were no significant differences between CD and WD males in tests for anxiety-like behavior, locomotor activity, spatial and object learning as well as social interest and social recognition in the Elevated-plus maze, light-dark box, Y-maze, NOR, and three-chamber test (**Suppl. Table 3**; **Suppl. Figure 1**).

#### Male attractiveness and reproductive tract parameters

Absolute testis weight was higher in WD compared to CD males (t = -2.956, p = 0.005; **Figure 3E**), however, relative testis weight (testis/body weight ratio) was lower (t = 3.045, p = 0.004). There was a trend for increased sperm numbers in WD males (t = -1.851, p = 0.077; **Figure 3F**), but lower testosterone levels (F_(1,2.511)_ = 49.831, p = 0.010; **Figure 3G**) compared to CD males. WD males had significantly lower total MUP concentration compared to CD males (t = 2.746, p = 0.025; **Figure 3H**), however, no relationship between body weight and MUP concentration or between body weight and protein/creatinine ratio was detected. Regarding testes morphological measures, no significant differences were detected (**Figure 3I-L**), but there was a trend for greater epithelium height in WD compared to CD males (t = 1.912, p = 0.092; **Figure 3J**).

#### Reproductive success, offspring sex ratio, and maternal care

There was no significant difference in reproductive success between groups as assessed by impregnated females or litter size (average litter sizes: WD = 6.1; CD = 6.5). However, the number of F1 males sired by WD males was significantly lower compared to CD males (t = 2.342, p = 0.025; **Figure 4D**). Maternal care differed between dams raising offspring from WD compared to CD males. The relative frequency of arched-back nursing (F_(1,18)_ = 6.270, p = 0.022; **Figure 4A**) and licking and grooming (F_(1,18)_ = 6.436, p = 0.021, **Figure 4B**) was lower in mothers of PatWD compared to mothers of PatCD offspring. Passive nursing occurred tentatively more often in mothers of PatWD compared to PatCD offspring (F_(1,19)_ = 3.459, p = 0.079, **Figure 4C**). The relative frequency of the dam not being on the nest was higher in mothers of PatWD compared to mothers of PatCD offspring (F_(1,18)_ = 4.611, p = 0.046). However, the total amount of nursing did not differ between groups. There was a trend for mothers of PatWD pups to have higher retrieving latencies than mothers of PatCD pups (t = 2.049; p = 0.080; **Figure 4E**).

**Figure 4.**
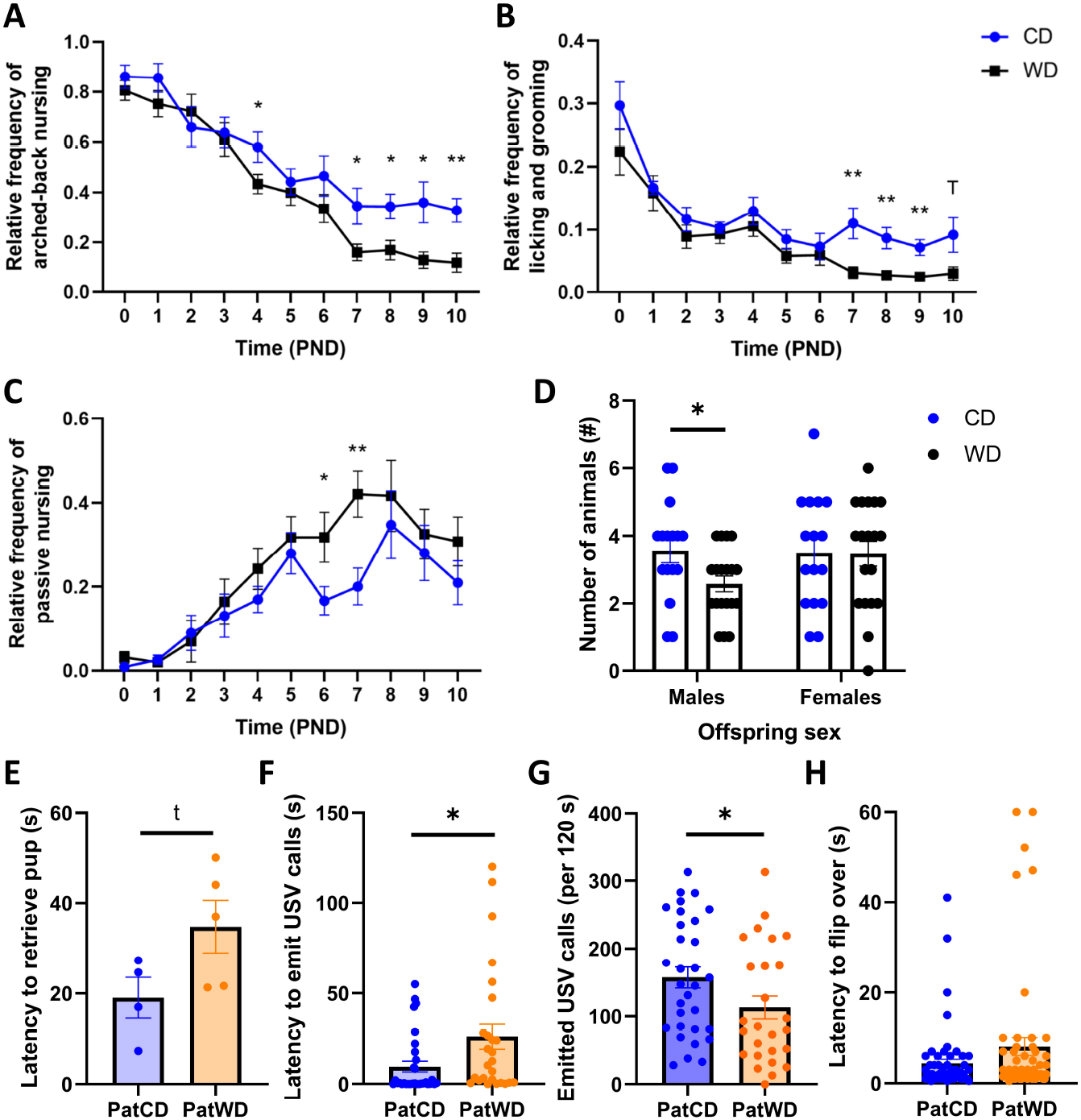
Maternal behavior of dams mated with F0 males, F1 offspring sex ratio, and F1 pup behavior. Maternal behavior as measured by **A)** arched-back nursing (N = 10-11), **B)** licking and grooming (N = 10-11), and **C)** passive nursing (N = 10-11). **D)** Offspring sex ratio (N = 16-19), **E)** mean latency of mothers to retrieve pups (N = 4-5), and pup behavior as reflected by **F)** latency to emit isolation-induced USV (N = 26-31) calls as well as **G)** number of USV calls (N = 26-31), and **H)** latency to flip over in surface righting test (N = 60-61). Graphs show group means ± SEM and, where possible, individual values. Statistics: General linear models (A-D), t-test (E-H). *, p < 0.05; **, p < 0.01.

### F1 generation

#### Neonatal testing

On PND 7, PatWD pups emitted USV calls after a significantly longer period of isolation from the mother (t = 2.868, p = 0.006; **Figure 4F**) and made significantly fewer calls (t = 2.278, p = 0.027) compared to PatCD pups (**Figure 4G**). The surface righting test did not reveal a difference between groups.

#### Physiologic parameters

Regarding body weight development (**Figure 5A**), the first body weight of F1 animals was assessed on PND 7. Taking into account the weekly body weights from week 1 to week 6, there was a significant effect of time (F_(2.6,528.3)_ = 7828.770, p < 0.001), a significant time x paternal diet interaction (F_(2.628,528.3)_ = 3.266, p = 0.026) and time x sex interaction (F_(2.628,528.3)_ = 125.705, p < 0.001). The analysis furthermore revealed a main effect of paternal diet (F_(1,201)_ = 7.540, p = 0.007), with PatWD mice weighing more than PatCD mice, and a significant main effect of sex (F_(1,201)_ = 71.403, p < 0.001), with males weighing more than females. *Post hoc* testing demonstrated that PatWD animals weighed significantly more compared to PatCD mice on PND 7 (F_(1,201)_ = 6.943, p = 0.009), 14 (F_(1,201)_ = 23.257, p < 0.001), and 21 (F_(1,201)_ = 20.924, p < 0.001; **Figure 5A**). From PND 28 on, PatCD and PatWD animals no longer differed in their body weights. Metabolic parameters measured by respiratory quotient **(Figure 5B)**, energy expenditure **(Figure 5C)**, food **(Figure 5D)** and water consumption **(Figure 5E)** as well as body composition assessed as fat mass **(Figure 5F)**, water **(Figure 5G)**, and lean mass **(Figure 5H)** did not differ between PatWD and PatCD mice. There was no significant effect of paternal diet on either baseline corticosterone or stress response levels (**Suppl. Table 4**).

**Figure 5.**
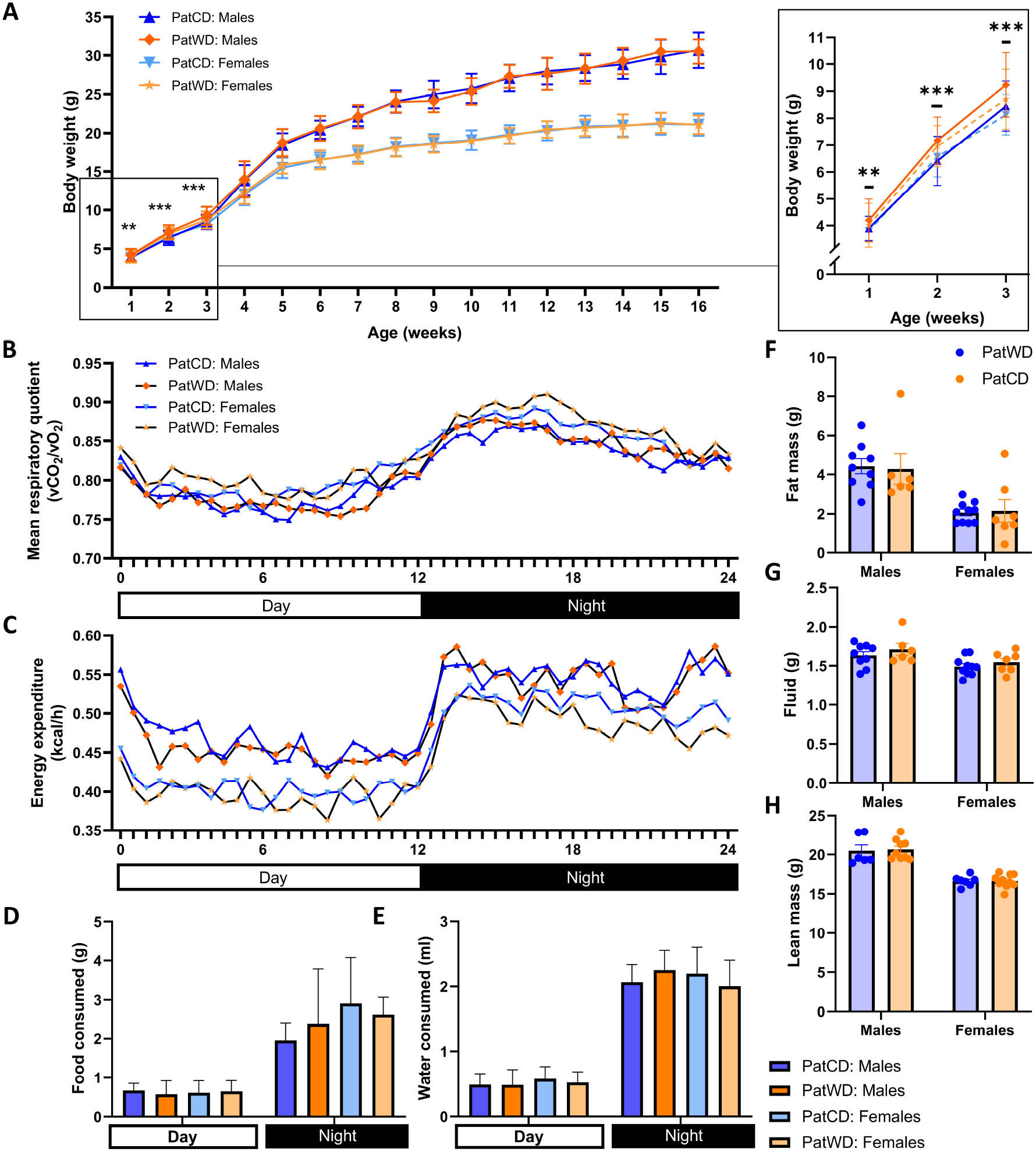
Body weights and metabolic parameters of F1 animals. **A)** Body weight development of male and female offspring, including a more detailed graph on the first three weeks. No differences were detected with regard to metabolic parameters gathered by the Promethion metabolic cage system, shown by **B**) mean respiratory quotient (N = 6-10) and **C)** mean energy expenditure during day and night (N = 6-10), **D)** food (N = 6-10), and **E)** water consumption (N = 6-10). Minispec measurements revealed that there were no differences in **F)** fat mass (N = 6-10), **G)** fluids (N = 6-10), and **H)** lean mass (N = 6-10) between PatWD and PatCD mice. Graphs show group means (± SEM) and individual values where possible. Statistics: General linear models, *post hoc* t-tests. *, p < 0.05; **, p < 0.01.

#### Gut microbiome

To evaluate the α-diversity of the microbiome in F1 mice, the Observed index (**Figure 6A**), Shannon’s diversity index (**Figure 6B**), and the inverse Simpson index (**Figure 6C**) were computed. No significant difference was found between paternal diet groups or sex. PCoA visualization of microbial β-diversity using the Bray–Curtis dissimilarity did not show clear clustering of the samples according to either paternal diet groups or sex (**Figure 6D**). PERMANOVA testing demonstrated that the microbial composition between the F1 mice of different paternal diet did not differ significantly. On the phylum level, ANCOM revealed that the abundance of Actinobacteria was significantly influenced by paternal diet, with PatWD mice having markedly lower proportions of Actinobacteria compared to PatCD mice (W-stat; **Figure 6E**). Further analysis revealed that *Actinobacteria* were mainly represented by the genus *Bifidobacterium* (**Figure 6F**), with their relative abundance being distinctly increased in PatWD compared to PatCD mice (W-stat; **Figure 6G**). No differences between groups were detected regarding other phyla or genuses.

**Figure 6.**
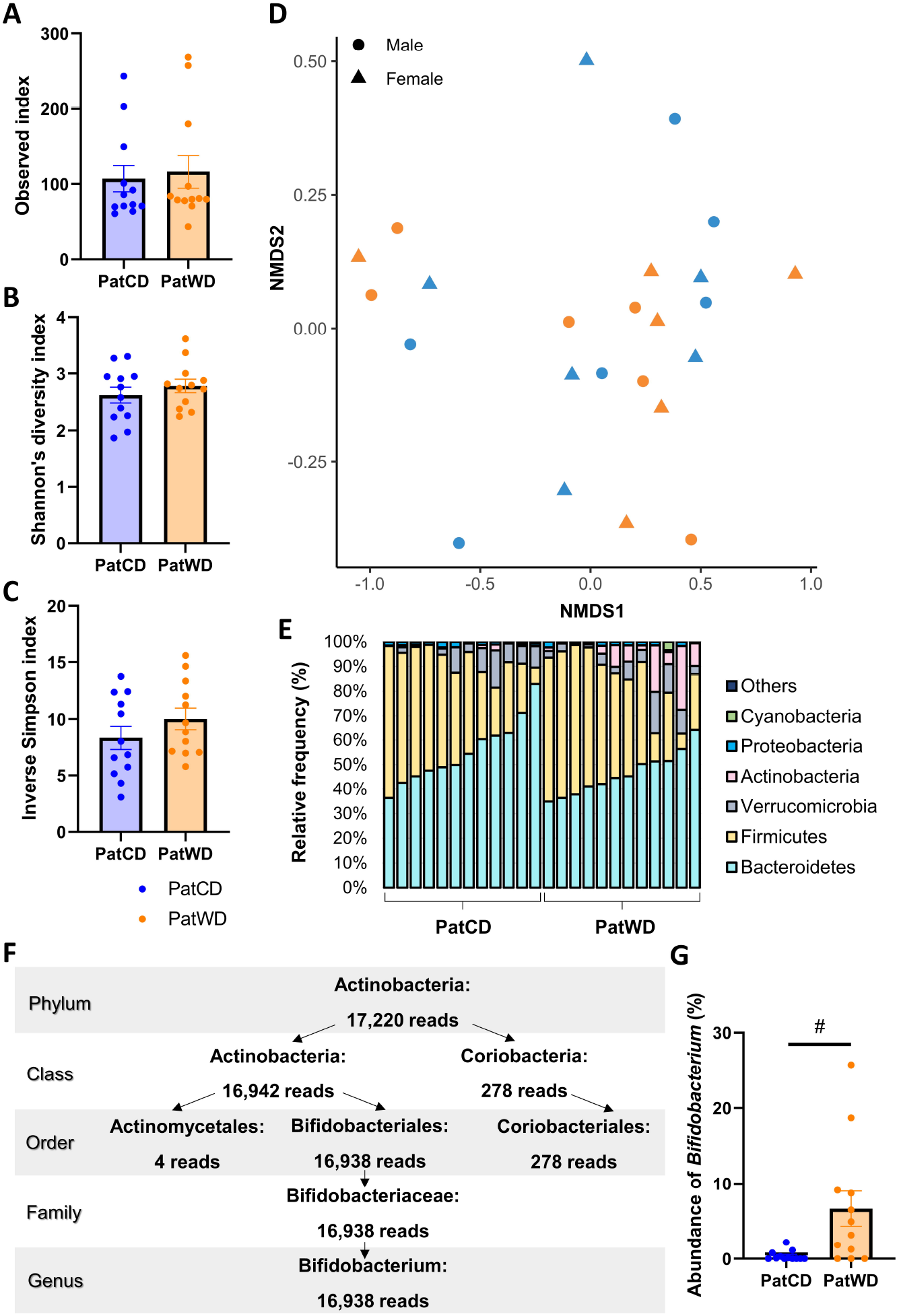
16S RNA analysis from faeces samples of F1 offspring. The α-diversity metric was calculated using the **A)** Observed index, **B)** Shannon’s diversity index, and the **C)** Inverse Simpson index and did not show significant differences between groups. **D)** NMDS plot demonstrating Bray-Curtis dissimilarity. **E)** ANCOM revealed a significant effect of paternal diet on *Actinobacteria* abundance at the phylum level. **F)** Further analysis of the *Actinobacteria* phylum showed that this phylum was mainly represented by the genus *Bifidobacterium*, with a **G)** distinctly increased relative abundance of *Bifidobacterium* in PatWD mice. Graphs show individual values and group means ± SEM. Statistics: t-test (A-C) or ANCOM (E-F); Sample size: N = 12. #, marked difference in ANCOM W-stat.

#### Behavioral tests

There were no significant differences between PatWD and PatCD regarding anxiety-like behavior (**Figure 7A-B**), exploratory locomotion (**Figure 7C**), risk assessment (**Figure 7D**), object recognition (**Figure 7F**), social interest (**Figure 7G**), social recognition (**Figure 7H**). With regard to spatial memory, as measured by the path covered and the time spent in the novel arm of the Y-maze, there was a significant difference regarding the distance in the novel arm (F_(1,47)_ = 4.225, p = 0.045) and a trend regarding the time in the novel arm (F_(1,47)_ = 3.654, p = 0.062; **Figure 7E**), with PatWD animals travelling significantly further in the novel arm and tending to spend more time in the novel arm compared to PatCD animals. The latency to immobility in the Porsolt swim test was significantly higher in PatWD compared to PatCD mice (F_(1,55)_ = 23.215, p < 0.001; **Figure 7I**), in both males (t = 3.423, p = 0.002) and females (t = 3.137, p = 0.006).

**Figure 7.**
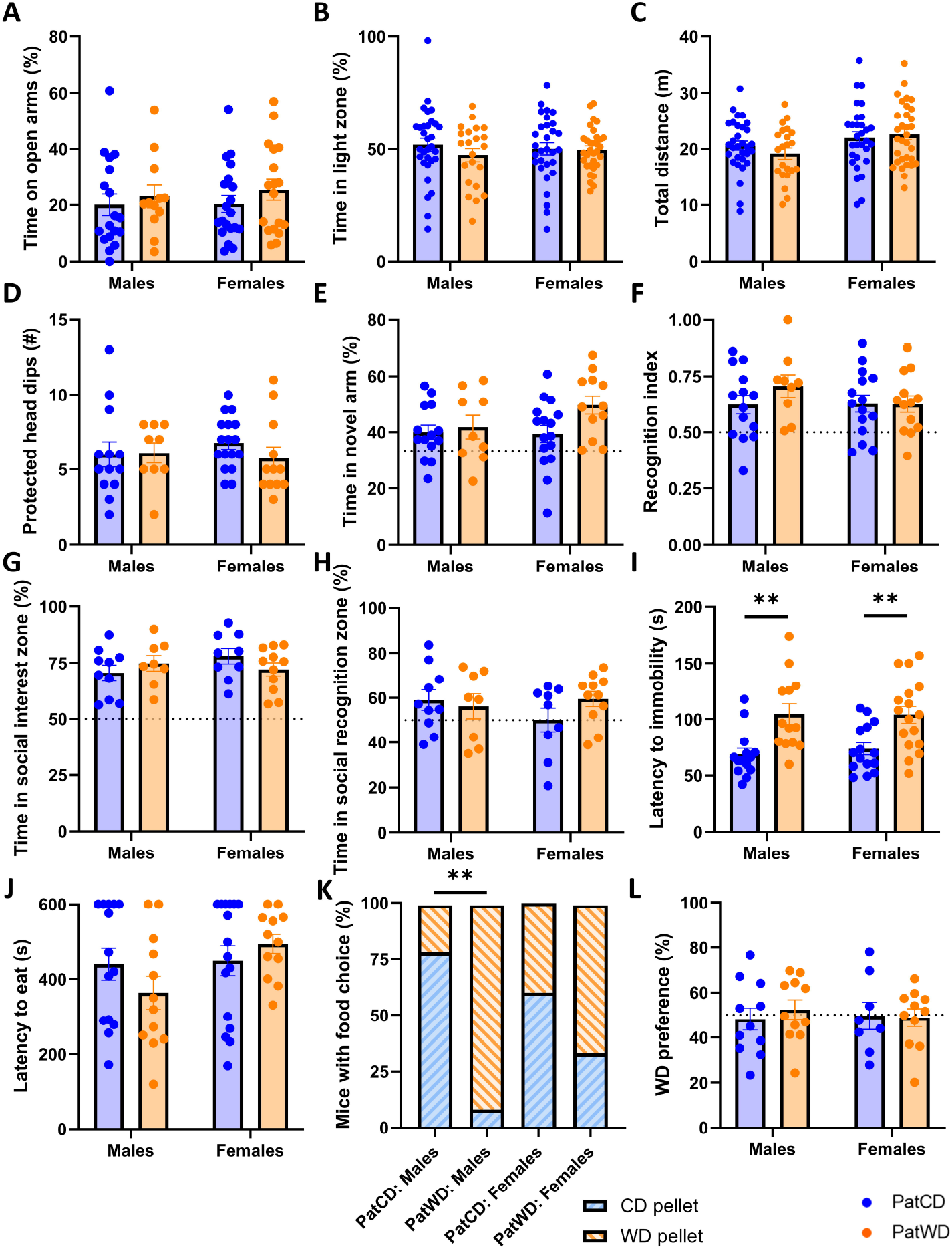
Anxiety, locomotion, learning, sociability, and food preference of F1 animals. No differences between PatWD and PatCD animals were detected regarding anxiety-like behavior as measured by **A)** time on open arms in the EPM (N = 12-20) and **B)** time spent in the light zone of the light-dark box (22-32) differed between PatWD and PatCD. Exploratory locomotion as reflected by the **C)** total distance travelled in the light-dark box (N = 22-32) and risk assessment assessed by **D)** protected head dips on the EPM (N = 9-16) did not differ between groups. There was a trend for PatWD mice to spend more **E)** time in the novel compared to all other arms in the Y-maze (9-16), reflecting spatial memory. Object memory was assessed by **F)** the recognition index in the novel object recognition (NOR) test (N = 9-16) and did not reveal a difference between groups. There were also no differences in **G)** sociability (N = 8-11) and **H)** social recognition in the three-chamber test (N = 8-11). **I)** The latency to become immobile in the Porsolt swim test differed significantly between paternal diet groups, with PatWD mice becoming immobile markedly later compared to PatCD mice (N = 13-17). **J)** Latency to start eating (N = 12-16) did not differ, but **K)** choice of food pellet (N = 12-16) differed between PatCD and PatWD mice in the novelty-suppressed feeding (NSF) paradigm. **L)** Smell preference of WD over CD pellet in a test based solely on olfactory cues of the pellets showed no difference between groups (N = 8-11). Graphs show individual values as well as group means ± SEM. Statistics: General linear models or χ2 test (K), *post hoc* t-test. **, p < 0.01.

To examine consummatory behavior, an NSF test was conducted where F1 mice were given a choice between an unfamiliar WD and a CD pellet. There was no difference in the latency to start eating (**Figure 7J**), however, the pellet choice significantly differed between male PatWD and male PatCD mice as significantly more PatWD mice started to eat the WD pellet compared to PatCD mice (X^2^ = 12.208, p < 0.001; **Figure 7K**). There was no difference between PatWD and PatCD mice in the food taste (both diets available ad libitum) and food smell preference tests (both diets presented in tea strainers; **Figure 7L**). Mice of both groups neither differed in the olfactory discrimination test (**Suppl. Figure 2A**) nor in the buried food test (**Suppl. Figure 2B**).

#### Parameters of mate attractiveness in male offspring

A significant main effect of paternal diet was found on male attractiveness, with PatWD males being preferred over PatCD males, in both the test with males and bedding (t = -5.563, p < 0.001; **Figure 8A**) and with bedding but without males (t = -5.597, p < 0.001; **Figure 8B**). PatWD males also won significantly more encounters in the dominance tube test (t = -5.134, p < 0.001; **Figure 8C**). While there was no difference in testes weight (**Figure 8D**) and sperm count (**Figure 8E**) between PatWD and PatCD males, they differed in their plasma testosterone levels. Specifically, PatWD males had lower levels of testosterone (t = -3.129, p = 0.006; **Figure 8F**). Neither total MUP concentration (**Figure 8G**) nor MUP/creatinine ratio (**Figure 8H**) differed between PatWD and PatCD males.

**Figure 8.**
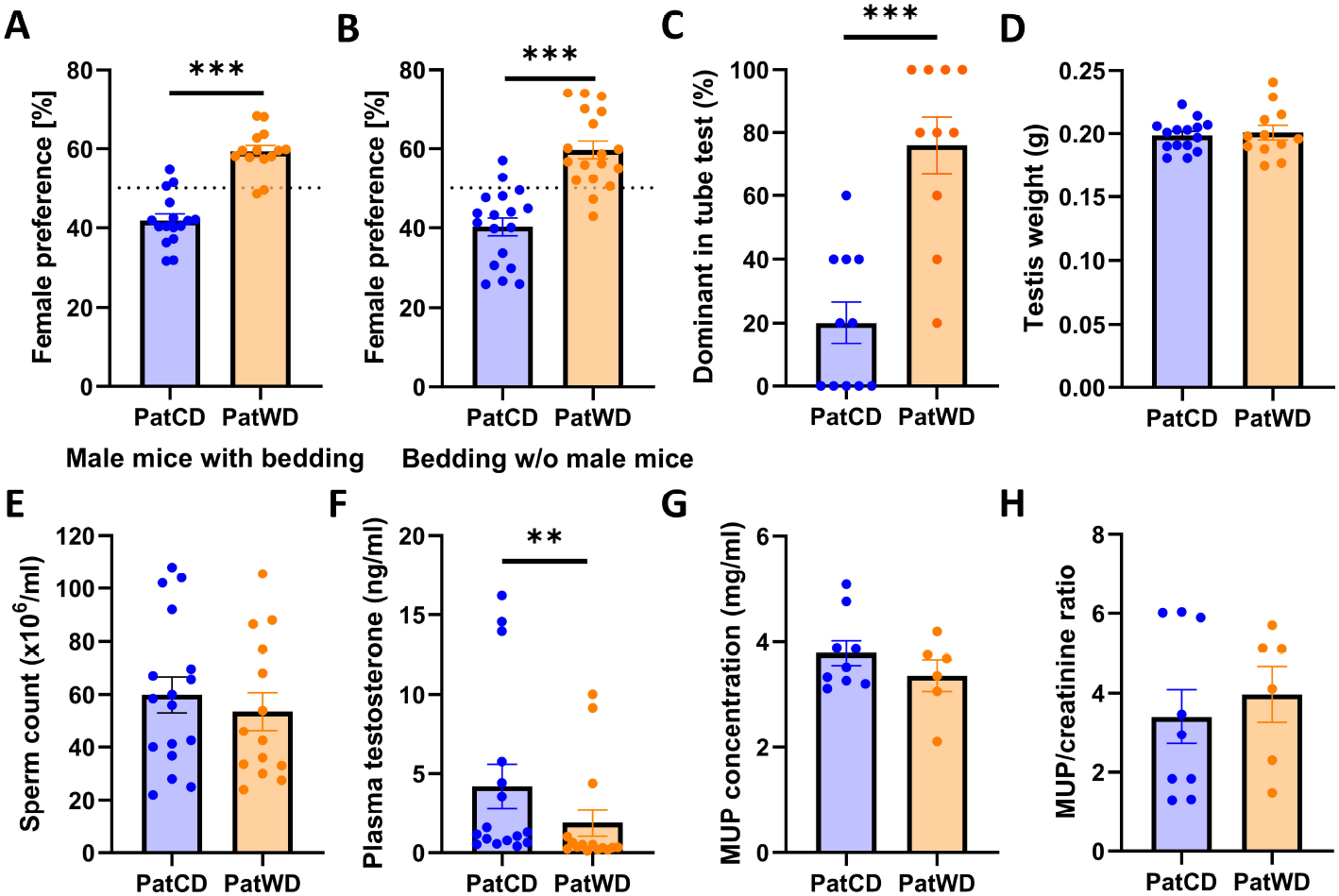
Female preference and parameters of reproductive fitness for F1 males. Female preference was assessed using **A)** both male mice and their soiled bedding in the three-chamber test (N = 14-16) and **B)** with male soiled bedding but without male mice (N = 14-16). Reproductive fitness was measured by means of **C)** the dominance tube test (N = 10-11), **D)** testis weight (N = 12-15), **E)** sperm count (N = 11-14), **F)** plasma testosterone levels (N = 15-16), **G)** total urinary MUP concentration (N = 6-9), and **H)** MUP/creatinine ratio (N = 6-9). Graphs show individual values and means ± SEM. Statistics: t-test. **, p < 0.01; ***, p < 0.001.

## Discussion

An unhealthy lifestyle may not only affect the individual, but also their progeny. It is becoming increasingly acknowledged that fathers can affect their offspring’s phenotype via epigenetic information, in addition to their genetic contribution ^8^. In this study, we investigated the influence of a Western-style diet (WD) during adolescence on male breeders and their offspring. By measuring both behavioral and physiological parameters in WD-fed males and their offspring, we were able to elucidate multiple aspects of paternal programming.

In F0 males, WD consumption during adolescence was reliably associated with increased body weight, food and calorie consumption, as well as increased fasting glucose levels compared to CD, which is in line with previous studies ^16, 31–36^. After only 6 weeks of WD consumption, mice showed metabolic changes indicative of a prediabetic state, reflected by the markedly increased body weight, fasting glucose levels, and fat pad mass with hypertrophied adipocytes. Histological analyses also confirmed that consumption of WD significantly increased liver mass, hepatic lipid droplet size and absolute testes mass, in accordance with previous studies ^33–35, 37–39^. In the present study, F0 mice were provided the WD only for 6 weeks, whereas other studies used longer periods of diet exposure and higher proportions of fat and/or sugar (e.g., ^10, 32^). As such, we show that already moderately increased amounts of fat and sugar over shorter periods of time during adolescence have significant effects on the body.

We also observed a decreased saccharin preference in WD compared to CD F0 males, a phenomenon that has been described previously (e.g., ^40^). The decreased saccharin preference following WD in F0 males could be explained by either anhedonia or by a devaluation of sweet tastes due to the abundance of sugar in the WD, which can lead to alterations or dysfunction of the brain’s reward systems ^41^. In contrast to previous studies ^37, 42, 43^, our results did not indicate increased anxiety-like behavior, decreased locomotor activity or impaired memory in WD compared to CD F0 males. The absence of these differences can likely be explained by lower fat and sugar proportions in the WD in comparison to other high-fat diets that used 60-80% fat, rapidly leading to obesity and impaired blood glucose control and insulin resistance, together with increased inactivity and learning impairments ^32, 35, 44^. However, it has recently been recommended to use a fat proportion between 30% and 50% when using a HFD as higher values could induce metabolic abnormalities ^15^. Interestingly, attractiveness towards females was decreased compared to control animals in WD males in the present study. Similar results have previously been reported for HFD-fed males ^45^. There is growing evidence that attractiveness is linked to the quantity of MUPs, with a higher concentration of MUPs acting as an attractant to females ^45^. MUPs are multiple proteins that are expressed in combinatorial patterns, differing between individuals, sex, and age ^46^. WD-fed F0 males in the present study had lower total MUP concentrations compared to CD males as a result of the diet intervention. This result corroborates previous findings that diet, such as HFD ^47^ or also dietary restriction ^48^, can influence MUP levels. As such, diet-induced differences in MUP levels most likely contributed to the observed difference in female preference.

Despite the decreased preference for WD males and less successful mountings compared to CD males, there was no difference in reproductive success, as reflected by percentage of pregnancies and offspring numbers. Hence, fertility was not significantly affected, which was confirmed by the sperm count that did not differ between experimental groups. Yet, there was a distinct difference in the sex ratio of offspring, with significantly fewer male offspring born to WD compared to CD fathers. It is currently unclear how and at what point in time (preconception, conception, prepartum) the sex ratio was potentially influenced, but there is increasing evidence that environmental factors, such as pollution and food scarcity, can modulate sex allocation in mammals ^9, 49–53^. In this study, a paternal WD resulted in female offspring favored over male offspring. Female offspring are generally considered more costly, and the sex ratio in mammals has been found to be skewed towards males under adverse environmental conditions and towards females under beneficial conditions ^51^. It was previously demonstrated that maternal diet, or more specifically, the proportion of saturated fats and carbohydrates in the diet of mothers, appears to be one cause for sex ratio distortion ^49, 50^. While traditionally the ability of fathers to bias offspring sex ratios has been dismissed, recent research has identified a male sperm morphological marker - the nucleus size – as a potential mechanism leading to sex ratio bias, explaining 22% variation in litter sex ratio ^53^. Future research could investigate whether the present sex-ratio bias associated with paternal WD can be traced back to sperm nucleus size.

There were also significant differences in maternal behavior towards offspring sired by WD or CD males. Mothers of PatCD offspring showed more high-quality maternal care, including arched-back nursing and licking and grooming, compared to mothers of PatWD offspring. This finding is in line with previously published data on decreased maternal care towards offspring from HFD compared to CD males ^37^. Korgan and colleagues ^37^ additionally found increased anxiety-like behavior in the offspring that received lower quality maternal care. It is well-acknowledged that variations in maternal care can cause behavioural changes in the offspring, especially regarding anxiety and fearfulness ^54^. The question remains why mothers of PatWD pups show lower quality maternal care compared to mothers of PatCD pups and whether this behavior reflects maternal programming. We furthermore detected fewer ultrasonic vocalization (USV) isolation calls emitted by PatWD offspring in comparison to PatCD offspring during the suckling period. Isolation calls are a common type of USVs that are produced by the pups upon isolation from their mother during the first two weeks of life ^55^. There might be a link between fewer pup vocalizations and lower quality of maternal care as it has been shown that the number of calls emitted by pups can reflect maternal responsiveness ^56^. Since mouse USV features are known to be highly consistent within the C57BL/6J strain, they represent a robust assay to test and demonstrate treatment effects ^55^. There is, however, a dispute as to whether pup isolation calls represent a negative affective state or whether they are simply the by-product of a thermoregulation process ^57^. Despite the differences in pup calls and maternal care, the total amount of nursing did not differ between groups. Nevertheless, there was a significant difference in offspring body weight, with PatWD mice weighing significantly more than PatCD mice on PND 7, 14, and 21, indicating that during early life when mice were housed with their mothers, PatWD pups had accelerated growth. Body weights of offspring no longer differed from PND 28 onward, when the young mice were weaned from their mothers. This observation might suggest that when a high energy food is available (i.e., the mother’s milk during nursing) the F1 PatWD offspring consumed more energy resulting in greater body weights compared to PatCD offspring. Future studies should examine how exposure to high energy diets during juvenile, adolescence and adult age groups physiologically impact offspring of PatWD and PatCD mice.

Gut microbiota analysis of F1 mice revealed a differential abundance of *Actinobacteria* in PatWD compared to PatCD animals on PND 28, with higher proportions of this phylum in PatWD mice. Previous research has shown that the relative proportion of the phylum *Actinobacteria* was elevated as a result of a HFD (45% fat) consumption in male mice and was positively associated with body weight and proinflammatory cytokines ^58^. In the present study, we demonstrate that the abundance of *Actinobacteria* was altered in the offspring of WD males. An increase in the relative proportion of *Actinobacteria* – as we found in PatWD offspring – has been linked to psychoactive or ‘psychobiotic’ effects, and to increased depressive-like behaviors ^59–61^. The phylum of *Actinobacteria* is mainly represented by the *Bifidobacterium* genus ^62^ – a finding that was supported by the present data. *Bifidobacterium* species are typically commensal bacteria that are known to have beneficial effects for health. *Bifidobacterium* abundance is typically decreased in diseases, however, increases have been reported in inflammatory bowel disease and allergies ^63, 64^. Given its role in gut health and immunomodulation, altered levels of *Bifidobacterium* could have influenced the offspring’s immune homeostasis. In humans, *Bifidobacterium longum* has been shown to be a modulator of stress responsiveness and cognition ^59^. As such, the present differences in Actinobacteria could have – in part – contributed to the F1 behavioral phenotype we observed, with enhanced spatial learning and decreased behavioral despair. While it is well acknowledged that mothers can influence their new-born’s gut microbiota (e.g. ^65^) less is known about the potential influence of the father ^66^. Here, we show that the relative abundance of the phylum of *Actinobacteria* in the offspring was associated with paternal diet, which could have major impacts on the offspring phenotype, as the microbiota can communicate with almost every other organ, including the brain ^67, 68^. Future studies are required to unravel potential mechanisms underpinning this change and whether a change of this phylum is causative for metabolic and/or psychological effects in the offspring.

Despite the observed changes in gut microbiota, there were no detectable differences in offspring metabolism, including energy expenditure, substrate oxidation (i.e., fatty acid and carbohydrate) and food intake. Furthermore, overall body composition did not differ between adult PatWD and PatCD mice. The finding that paternal diet did not impact metabolism is important, because previous studies have shown marked effects of paternal diet on offspring body weight development and metabolism ^17, 35, 37, 69^. The differences between this and previous studies might be explained by differences in diet composition, where we employed a 21% saturated fat WD for 6 weeks compared to others that employed more extreme diets consisting of >60% fat content and/or longer durations of up to 6 months ^17, 32, 35^. The diet employed herein more accurately represents a common WD and for this reason our findings are of broad translational relevance ^70^. It is also possible that a ‘second hit’ ^71, 72^, such as a genetic or environmental factor, might be required to induce metabolic consequences for PatWD compared to PatCD offspring.

Behavioral testing of offspring revealed no difference in anxiety-like behavior, exploratory locomotion, spatial learning, object learning, social interaction and social recognition, and risk assessment between PatWD and PatCD animals. However, the latency to immobility in the Porsolt test was significantly longer in PatWD animals compared to PatCD mice, suggesting reduced susceptibility to behavioral despair, or resilience to stress in this environment. As opposed to studies proposing that a maternal unhealthy diet promotes anxiogenic and depression-like behaviors in the progeny ^73, 74^, a paternal WD therefore rather predicts reduced behavioural despair. These findings stand in contrast to observations in prior paternal HFD studies.

In order to elucidate a potentially inherited preference of PatWD mice for WD, several tests were performed that assessed food preference based on taste and smell or smell alone. The finding that PatWD males initially chose to eat the WD over the CD pellet in the NSF, while PatCD males initially preferred to eat the CD pellet, is indicative of adaptive paternal programming. The follow-up experiments examined olfactory impairments or lack of motivation to eat and showed no differences between groups. However, we demonstrated that the food preference was absent when mice could only perceive olfactory cues but were not allowed to lick and taste them. The senses of both taste and smell play a key role in the sensory effects on food choice and intake ^75^. Here, only taste, but not smell alone was linked to a preference, and thus seemed to be a rewarding stimulus. While the sense of smell has a priming role in eating behavior, taste plays a role as (macro)nutrient sensing system during consumption. It has been shown that odor exposure can induce appetite specifically for the cued food. If and how odors modulate food choice and intake is less clear ^75^.

The female preference test revealed both a preference for PatWD over PatCD males, and their soiled bedding, respectively. This preference cannot be explained by commonly discussed attractiveness parameters, such as the body size of the males or their testes (which did not differ between groups), or higher testosterone levels, as PatWD animals had significantly lower values compared to PatCD males. However, the dominance tube test revealed that PatWD males were dominant over PatCD males, demonstrating a behavioral phenotype of dominance in PatWD males, despite no body weight differences. Previously, it has been shown that male major urinary protein (MUP) levels are associated with social status in mouse social hierarchies, with dominant males producing more MUPs than subordinates ^76^. However, we did not detect differential MUP concentrations between PatWD and PatCD mice.

Altogether, several distinct phenotypic effects were observed, resulting from either acute or paternal WD. While we could not recapitulate some of the phenotypic observations made in prior studies using higher proportions of fat or sugar (e.g., ^10, 32^), the present study aimed for a comprehensive picture of the intergenerational effects of paternal WD. The different outcomes are most likely the result of multifactorial causes, including different proportions of fat and sugar in the diet, the duration of the diet manipulation, the timing of mating, whether *in vitro* fertilisation was performed or natural conception occurred, the species, strain, and origin of the animals, and the timing and performance of the respective test itself, relating to the current debate about the ‘reproducibility crisis’ and going far beyond the scope of this article ^77^. However, future experiments with higher proportions of fat and/or sugar in the paternal diet would be interesting to find out whether endophenotypes are exacerbated in this way.

## Conclusions

In summary, a Western-style diet consumed across puberty and adolescence was associated with increased F0 body weight, decreased attractiveness of F0 male mice, and lower quality of maternal care. Furthermore, we discovered striking intergenerational impacts of paternal WD, including higher body weights of F1 pups during the suckling period, increased *Actinobacteria* abundance in the gut, increased attractiveness, and dominance of male offspring, and decreased behavioral despair. These findings indicate a potential ‘resiliency’ phenotype in the PatWD offspring. Our observations seemingly suggest beneficial effects of a paternal WD on male F1 offspring reproductive parameters, although this specific change in the gut microbiota could have deleterious consequences. A paternally mediated intergenerational impact on gut microbiota has not been described and could have major impacts on the offspring phenotype. In light of the evidence that WD and related dietary exposures can alter human sperm epigenetics, and that gut microbiota changes have been associated with many different disorders, the present findings have significant public health implications.

## Declarations

### Ethics approval

All experimental procedures were conducted following the *Australian code for the care and use of animals for scientific purposes* (National Health and Medical Research Council [NHMRC], 2013). The experiments were approved by the Animal Ethics Committee of the Florey Institute of Neuroscience and Mental Health.

### Availability of data

The datasets used and/or analyzed during the current study are available from the corresponding author on reasonable request.

## Acknowledgements

The Florey Institute of Neuroscience and Mental Health acknowledges the strong support from the Victorian Government and in particular the funding from the Operational Infrastructure Support Grant. This work was supported by infrastructure from the Melbourne Mouse Metabolic Phenotyping Platform (MMMPP) at the University of Melbourne and excellent technical expertise of Vanessa Haynes.

## Authors’ contributions

CB: Conceptualization; Data curation; Formal analysis; Funding acquisition; Investigation; Methodology; Project administration; Resources; Software; Supervision; Validation; Visualization; Writing - original draft; Writing - review & editing. TP: Data curation; Investigation; Methodology; Resources; Writing - review & editing. YF: Data curation; Investigation; Methodology; Writing - review & editing. FM: Data curation; Investigation; Methodology; SL: Data curation; Investigation; Methodology; Writing - review & editing. GK: Methodology; Software; Validation; Writing - review & editing. MJW: Conceptualization; Methodology; Project administration; Resources; Writing - review & editing. ACR: Conceptualization; Funding acquisition; Project administration; Supervision; Writing - review & editing. AJH: Conceptualization; Funding acquisition; Methodology; Project administration; Resources; Supervision; Writing - review & editing.

## Nonstandard abbreviations

CD: Control diet (7% fat, 10% sugar)
DOHaD: Developmental Origins of Health and Disease
F0: Parental generation
F1: Filial generation
FFT: Fast Fourier transformation
GTT: Glucose tolerance test
H&E: Hematoxylin and eosin
MUPs: Major urinary proteins
NOR: Novel object recognition
NSF: Novelty-suppressed feeding
PatCD: Paternal CD, i.e., the father was fed the control diet preconception
PatWD: Paternal WD, i.e., the father was fed the Western diet preconception
PND: Postnatal day
USV: Ultrasonic vocalization
WD: Western-style high-fat (21%) and high-sugar (34%) diet

## References

1. World Health Organization Diet, nutrition and the prevention of chronic diseases: report of a Joint WHO/FAO: Geneva: World Health Organization 2003: Geneva: World Health Organization.

2. Bellisari A Evolutionary origins of obesity. Obes. Rev. 2008;9:165–180.

3. World Health Organization 2020 Obesity and overweight factsheet. https://www.who.int/news-room/fact-sheets/detail/obesity-and-overweight. Accessed May 2020.

4. NCHS 1999-2016 National health and nutrition examination survey (NHANES) 1999-2016. Centers for Disease Control and Prevention (CDC). National Center for Health Statistics (NCHS):Hyattsville, MD: U.S. Department of Health and Human Services, Centers for Disease Control and Prevention. https://www.cdc.gov/nchs/data/nhanes/.

5. Haslam DW, James WPT Obesity. Lancet 2005;366:1197–1209.

6. Wiklund P The role of physical activity and exercise in obesity and weight management: Time for critical appraisal. J. Sport Health Sci. 2016;5:151–154.

7. Lohse T, Rohrmann S, Bopp M, Faeh D Heavy Smoking Is More Strongly Associated with General Unhealthy Lifestyle than Obesity and Underweight. PloS one 2016;11:e0148563.

8. Bodden C, Hannan AJ, Reichelt AC Diet-Induced Modification of the Sperm Epigenome Programs Metabolism and Behavior. Trends Endocrinol. Metab. 2020;31:131–149.

9. Edwards AM, Cameron EZ Forgotten fathers: paternal influences on mammalian sex allocation. Trends in ecology & evolution 2014;29:158–164.

10. Huypens P, Sass S, Wu M, Dyckhoff D, Tschöp M, Theis F, Marschall S, Hrabě de Angelis M, Beckers J Epigenetic germline inheritance of diet-induced obesity and insulin resistance. Nat. Genet. 2016;48:497–499.

11. Lowe CJ, Morton JB, Reichelt AC Adolescent obesity and dietary decision making—a brain-health perspective. Lancet Child Adolesc. Health 2020;4:388–396.

12. Nätt D, Kugelberg U, Casas E, Nedstrand E, Zalavary S, Henriksson P, Nijm C, Jäderquist J, Sandborg J, Flinke E, Ramesh R, Örkenby L, Appelkvist F, Lingg T, Guzzi N, Bellodi C, Löf M, Vavouri T, Öst A Human sperm displays rapid responses to diet. PLoS biology 2019;17:e3000559.

13. Ibáñez CA, Erthal RP, Ogo FM, Peres MNC, Vieira HR, Conejo C, Tófolo LP, Francisco FA, da Silva Silveira S, Malta A, Pavanello A, Martins IP, da Silva Paulo H. O., Jacinto Saavedra LP, Gonçalves GD, Moreira VM, Alves VS, da Silva Franco, Claudinéia C., Previate C, Gomes RM, Oliveira Venci R de, Dias FRS, Armitage JA, Zambrano E, Mathias PCF, Fernandes GSA, Palma-Rigo K A High Fat Diet during Adolescence in Male Rats Negatively Programs Reproductive and Metabolic Function Which Is Partially Ameliorated by Exercise. Front. Physiol. 2017;8:807.

14. Laviola G, Macrı S, Morley-Fletcher S, Adriani W Risk-taking behavior in adolescent mice: psychobiological determinants and early epigenetic influence. Neurosci. Biobehav. Rev. 2003;27:19–31.

15. Rodríguez-Correa E, González-Pérez I, Clavel-Pérez PI, Contreras-Vargas Y, Carvajal K Biochemical and nutritional overview of diet-induced metabolic syndrome models in rats: what is the best choice? Nutr. Diabetes 2020;10:24.

16. McPherson NO, Fullston T, Bakos HW, Setchell BP, Lane M Obese father’s metabolic state, adiposity, and reproductive capacity indicate son’s reproductive health. Fertility and sterility 2014;101:865–873.

17. Fullston T, Ohlsson Teague EMC, Palmer NO, DeBlasio MJ, Mitchell M, Corbett M, Print CG, Owens JA, Lane M Paternal obesity initiates metabolic disturbances in two generations of mice with incomplete penetrance to the F2 generation and alters the transcriptional profile of testis and sperm microRNA content. FASEB J. 2013;27:4226–4243.

18. Fennell KA, Busby RGG, Li S, Bodden C, Stanger SJ, Nixon B, Short AK, Hannan AJ, Pang TY Limitations to intergenerational inheritance: subchronic paternal stress preconception does not influence offspring anxiety. Scientific reports 2020;10:16050.

19. Ribeiro CT, Souza DB de, Costa WS, Sampaio FJB, Pereira-Sampaio MA Immediate and late effects of chronic stress in the testes of prepubertal and adult rats. Asian J. Androl. 2018;20:385–390.

20. Beynon RJ, Veggerby C, Payne CE, Robertson DHL, Gaskell SJ, Humphries RE, Hurst JL Polymorphism in major urinary proteins: molecular heterogeneity in a wild mouse population. J. Chem. Ecol. 2002;28:1429–1446.

21. Garratt M, Stockley P, Armstrong SD, Beynon RJ, Hurst JL The scent of senescence: sexual signalling and female preference in house mice. Journal of evolutionary biology 2011;24:2398–2409.

22. Thompson LR, Sanders JG, McDonald D, Amir A, Ladau J, Locey KJ, Prill RJ, Tripathi A, Gibbons SM, Ackermann G, Navas-Molina JA, Janssen S, Kopylova E, Vázquez-Baeza Y, González A, Morton JT, Mirarab S, Zech Xu Z, Jiang L, Haroon MF, Kanbar J, Zhu Q, Jin Song S, Kosciolek T, Bokulich NA, Lefler J, Brislawn CJ, Humphrey G, Owens SM, Hampton-Marcell J, Berg-Lyons D, McKenzie V, Fierer N, Fuhrman JA, Clauset A, Stevens RL, Shade A, Pollard KS, Goodwin KD, Jansson JK, Gilbert JA, Knight R A communal catalogue reveals Earth’s multiscale microbial diversity. Nature 2017;551:457–463.

23. Gonzalez A, Navas-Molina JA, Kosciolek T, McDonald D, Vázquez-Baeza Y, Ackermann G, DeReus J, Janssen S, Swafford AD, Orchanian SB, Sanders JG, Shorenstein J, Holste H, Petrus S, Robbins-Pianka A, Brislawn CJ, Wang M, Rideout JR, Bolyen E, Dillon M, Caporaso JG, Dorrestein PC, Knight R Qiita: rapid, web-enabled microbiome meta-analysis. Nat Methods 2018;15:796–798.

24. Bolyen E, Rideout JR, Dillon MR, Bokulich NA, Abnet CC, Al-Ghalith GA, Alexander H, Alm EJ, Arumugam M, Asnicar F, Bai Y, Bisanz JE, Bittinger K, Brejnrod A, Brislawn CJ, Brown CT, Callahan BJ, Caraballo-Rodríguez AM, Chase J, Cope EK, Da Silva R, Diener C, Dorrestein PC, Douglas GM, Durall DM, Duvallet C, Edwardson CF, Ernst M, Estaki M, Fouquier J, Gauglitz JM, Gibbons SM, Gibson DL, Gonzalez A, Gorlick K, Guo J, Hillmann B, Holmes S, Holste H, Huttenhower C, Huttley GA, Janssen S, Jarmusch AK, Jiang L, Kaehler BD, Kang KB, Keefe CR, Keim P, Kelley ST, Knights D, Koester I, Kosciolek T, Kreps J, Langille MGI, Lee J, Ley R, Liu Y-X, Loftfield E, Lozupone C, Maher M, Marotz C, Martin BD, McDonald D, McIver LJ, Melnik AV, Metcalf JL, Morgan SC, Morton JT, Naimey AT, Navas-Molina JA, Nothias LF, Orchanian SB, Pearson T, Peoples SL, Petras D, Preuss ML, Pruesse E, Rasmussen LB, Rivers A, Robeson MS, Rosenthal P, Segata N, Shaffer M, Shiffer A, Sinha R, Song SJ, Spear JR, Swafford AD, Thompson LR, Torres PJ, Trinh P, Tripathi A, Turnbaugh PJ, Ul-Hasan S, van der Hooft JJJ, Vargas F, Vázquez-Baeza Y, Vogtmann E, Hippel M von, Walters W, Wan Y, Wang M, Warren J, Weber KC, Williamson CHD, Willis AD, Xu ZZ, Zaneveld JR, Zhang Y, Zhu Q, Knight R, Caporaso JG Reproducible, interactive, scalable and extensible microbiome data science using QIIME 2. Nat Biotechnol 2019;37:852–857.

25. Short AK, Fennell KA, Perreau VM, Fox A, O’Bryan MK, Kim JH, Bredy TW, Pang TY, Hannan AJ Elevated paternal glucocorticoid exposure alters the small noncoding RNA profile in sperm and modifies anxiety and depressive phenotypes in the offspring. Translational psychiatry 2016;6:e837.

26. Leger M, Quiedeville A, Bouet V, Haelewyn B, Boulouard M, Schumann-Bard P, Freret T Object recognition test in mice. Nat Protoc 2013;8:2531–2537.

27. Screven LA, Dent ML Preference in female laboratory mice is influenced by social experience. Behavioural processes 2018;157:171–179.

28. Mitra R, Sapolsky RM Short-term enrichment makes male rats more attractive, more defensive and alters hypothalamic neurons. PLoS ONE 2012;7:e36092.

29. Faul F, Erdfelder E, Lang A-G, Buchner A G*Power 3: A flexible statistical power analysis program for the social, behavioral, and biomedical sciences. Behav. Res. Methods 2007;39:175–191.

30. Cohen J Statistical power analysis for the behavioral sciences: New York. Routledge 1988: New York.

31. Castro Barbosa T de, Ingerslev LR, Alm PS, Versteyhe S, Massart J, Rasmussen M, Donkin I, Sjögren R, Mudry JM, Vetterli L, Gupta S, Krook A, Zierath JR, Barrès R High-fat diet reprograms the epigenome of rat spermatozoa and transgenerationally affects metabolism of the offspring. Molecular metabolism 2016;5:184–197.

32. Chen Q, Yan M, Cao Z, Li X, Zhang Y, Shi J, Feng G, Peng H, Zhang X, Zhang Y, Qian J, Duan E, Zhai Q, Zhou Q Sperm tsRNAs contribute to intergenerational inheritance of an acquired metabolic disorder. Science (New York, N.Y.) 2016;351:397–400.

33. Fan Y, Liu Y, Xue K, Gu G, Fan W, Xu Y, Ding Z Diet-induced obesity in male C57BL/6 mice decreases fertility as a consequence of disrupted blood-testis barrier. PloS one 2015;10:e0120775.

34. Ferramosca A, Conte A, Moscatelli N, Zara V A high-fat diet negatively affects rat sperm mitochondrial respiration. Andrology 2016;4:520–525.

35. Fontelles CC, Guido LN, Rosim MP, Andrade FdO, Jin L, Inchauspe J, Pires VC, Castro IA de, Hilakivi-Clarke L, Assis S de, Ong TP Paternal programming of breast cancer risk in daughters in a rat model: opposing effects of animal-and plant-based high-fat diets. Breast Cancer Res. 2016;18.

36. Ng S-F, Lin RCY, Laybutt DR, Barrès R, Owens JA, Morris MJ Chronic high-fat diet in fathers programs β-cell dysfunction in female rat offspring. Nature 2010;467:963–966.

37. Korgan AC, O’Leary E, King JL, Weaver ICG, Perrot TS Effects of paternal high-fat diet and rearing environment on maternal investment and development of defensive responses in the offspring. Psychoneuroendocrinology 2018;91:20–30.

38. Watkins AJ, Dias I, Tsuro H, Allen D, Emes RD, Moreton J, Wilson R, Ingram RJM, Sinclair KD Paternal diet programs offspring health through sperm-and seminal plasma-specific pathways in mice. PNAS 2018;115:10064–10069.

39. Mitchell M, Bakos HW, Lane M Paternal diet-induced obesity impairs embryo development and implantation in the mouse. Fertility and sterility 2011;95:1349–1353.

40. Rabasa C, Winsa-Jörnulf J, Vogel H, Babaei CS, Askevik K, Dickson SL Behavioral consequences of exposure to a high fat diet during the post-weaning period in rats. Horm. Behav. 2016;85:56–66.

41. Peciña S, Cagniard B, Berridge KC, Aldridge JW, Zhuang X Hyperdopaminergic Mutant Mice Have Higher “Wanting” But Not “Liking” for Sweet Rewards. J. Neurosci. 2003;23:9395–9402.

42. Sharma S, Fulton S Diet-induced obesity promotes depressive-like behaviour that is associated with neural adaptations in brain reward circuitry. International journal of obesity (2005) 2013;37:382–389.

43. Hassan AM, Mancano G, Kashofer K, Fröhlich EE, Matak A, Mayerhofer R, Reichmann F, Olivares M, Neyrinck AM, Delzenne NM, Claus SP, Holzer P High-fat diet induces depression-like behaviour in mice associated with changes in microbiome, neuropeptide Y, and brain metabolome. Nutritional Neuroscience 2019;22:877–893.

44. An T, Zhang T, Teng F, Zuo J-C, Pan Y-Y, Liu Y-F, Miao J-N, Gu Y-J, Yu N, Zhao D-D, Mo F-F, Gao S-H, Jiang G Long non-coding RNAs could act as vectors for paternal heredity of high fat diet-induced obesity. Oncotarget 2017;8:47876–47889.

45. Kumar V, Vasudevan A, Soh LJT, Le Min C, Vyas A, Zewail-Foote M, Guarraci FA Sexual attractiveness in male rats is associated with greater concentration of major urinary proteins. Biology of reproduction 2014;91:150.

46. Logan DW, Marton TF, Stowers L Species Specificity in Major Urinary Proteins by Parallel Evolution. PLoS ONE 2008;3.

47. Kleinert M, Parker BL, Jensen TE, Raun SH, Pham P, Han X, James DE, Richter EA, Sylow L Quantitative proteomic characterization of cellular pathways associated with altered insulin sensitivity in skeletal muscle following high-fat diet feeding and exercise training. Sci Rep 2018;8:10723.

48. Giller K, Huebbe P, Doering F, Pallauf K, Rimbach G Major urinary protein 5, a scent communication protein, is regulated by dietary restriction and subsequent re-feeding in mice. Proceedings. Biological sciences 2013;280:20130101.

49. Rosenfeld CS, Grimm KM, Livingston KA, Brokman AM, Lamberson WE, Roberts RM Striking variation in the sex ratio of pups born to mice according to whether maternal diet is high in fat or carbohydrate. Proceedings of the National Academy of Sciences of the United States of America 2003;100:4628–4632.

50. Rosenfeld CS, Roberts RM Maternal diet and other factors affecting offspring sex ratio: a review. Biology of reproduction 2004;71:1063–1070.

51. Schacht R, Tharp D, Smith KR Sex ratios at birth vary with environmental harshness but not maternal condition. Scientific reports 2019;9:9066.

52. Öst A, Lempradl A, Casas E, Weigert M, Tiko T, Deniz M, Pantano L, Boenisch U, Itskov PM, Stoeckius M, Ruf M, Rajewsky N, Reuter G, Iovino N, Ribeiro C, Alenius M, Heyne S, Vavouri T, Pospisilik JA Paternal diet defines offspring chromatin state and intergenerational obesity. Cell 2014;159:1352–1364.

53. Malo AF, Martinez-Pastor F, Garcia-Gonzalez F, Garde J, Ballou JD, Lacy RC A father effect explains sex-ratio bias. Proceedings. Biological sciences 2017;284.

54. Caldji C, Tannenbaum B, Sharma S, Francis D, Plotsky PM, Meaney MJ Maternal care during infancy regulates the development of neural systems mediating the expression of fearfulness in the rat. Proc. Natl. Acad. Sci. U.S.A. 1998;95:5335–5340.

55. Barnes TD, Rieger MA, Dougherty JD, Holy TE Group and Individual Variability in Mouse Pup Isolation Calls Recorded on the Same Day Show Stability. Front. Behav. Neurosci. 2017;11:243.

56. D’Amato FR, Scalera E, Sarli C, Moles A Pups call, mothers rush: does maternal responsiveness affect the amount of ultrasonic vocalizations in mouse pups? Behav. Genet. 2005;35:103–112.

57. Blumberg MS, Sokoloff G Do infant rats cry? Psychol. Rev. 2001;108:83–95.

58. Kim SJ, Kim S-E, Kim A-R, Kang S, Park M-Y, Sung M-K Dietary fat intake and age modulate the composition of the gut microbiota and colonic inflammation in C57BL/6J mice. BMC Microbiol 2019;19:193.

59. Allen AP, Hutch W, Borre YE, Kennedy PJ, Temko A, Boylan G, Murphy E, Cryan JF, Dinan TG, Clarke G Bifidobacterium longum 1714 as a translational psychobiotic: modulation of stress, electrophysiology and neurocognition in healthy volunteers. Transl Psychiatry 2016;6:e939.

60. Zheng P, Zeng B, Zhou C, Liu M, Fang Z, Xu X, Zeng L, Chen J, Fan S, Du X, Zhang X, Yang D, Yang Y, Meng H, Li W, Melgiri ND, Licinio J, Wei H, Xie P Gut microbiome remodeling induces depressive-like behaviors through a pathway mediated by the host’s metabolism. Mol Psychiatry 2016;21:786–796.

61. Barandouzi ZA, Starkweather AR, Henderson WA, Gyamfi A, Cong XS Altered Composition of Gut Microbiota in Depression: A Systematic Review. Front. Psychiatry 2020;11:541.

62. Rinninella E, Raoul P, Cintoni M, Franceschi F, Miggiano GAD, Gasbarrini A, Mele MC What is the Healthy Gut Microbiota Composition? A Changing Ecosystem across Age, Environment, Diet, and Diseases. Microorganisms 2019;7.

63. Wang W, Chen L, Zhou R, Wang X, Song L, Huang S, Wang G, Xia B Increased proportions of Bifidobacterium and the Lactobacillus group and loss of butyrate-producing bacteria in inflammatory bowel disease. Journal of Clinical Microbiology 2014;52:398–406.

64. Simonyté Sjödin K, Hammarström M-L, Rydén P, Sjödin A, Hernell O, Engstrand L, West CE Temporal and long-term gut microbiota variation in allergic disease: A prospective study from infancy to school age. Allergy 2019;74:176–185.

65. Macpherson AJ, Agüero MG de, Ganal-Vonarburg SC How nutrition and the maternal microbiota shape the neonatal immune system. Nature reviews. Immunology 2017;17:508–517.

66. Bromfield JJ, Schjenken JE, Chin PY, Care AS, Jasper MJ, Robertson SA Maternal tract factors contribute to paternal seminal fluid impact on metabolic phenotype in offspring. PNAS 2014;111:2200–2205.

67. Collins SM, Surette M, Bercik P The interplay between the intestinal microbiota and the brain. Nat Rev Microbiol 2012;10:735–742.

68. Cryan JF, Dinan TG Mind-altering microorganisms: the impact of the gut microbiota on brain and behaviour. Nature reviews. Neuroscience 2012;13:701–712.

69. Consitt LA, Saxena G, Slyvka Y, Clark BC, Friedlander M, Zhang Y, Nowak FV Paternal high-fat diet enhances offspring whole-body insulin sensitivity and skeletal muscle insulin signaling early in life. Physiological reports 2018;6.

70. Speakman JR Use of high-fat diets to study rodent obesity as a model of human obesity. Int J Obes 2019;43:1491–1492.

71. Bayer TA, Falkai P, Maier W Genetic and non-genetic vulnerability factors in schizophrenia: the basis of the “Two hit hypothesis”. J. Psychiatr. Res. 1999;33:543–548.

72. Walker AK, Nakamura T, Byrne RJ, Naicker S, Tynan RJ, Hunter M, Hodgson DM Neonatal lipopolysaccharide and adult stress exposure predisposes rats to anxiety-like behaviour and blunted corticosterone responses: implications for the double-hit hypothesis. Psychoneuroendocrinology 2009;34:1515–1525.

73. Winther G, Elfving B, Müller HK, Lund S, Wegener G Maternal High-fat Diet Programs Offspring Emotional Behavior in Adulthood. Neuroscience 2018;388:87–101.

74. Bergmann S, Schlesier-Michel A, Wendt V, Grube M, Keitel-Korndörfer A, Gausche R, Klitzing K von, Klein AM Maternal Weight Predicts Children’s Psychosocial Development via Parenting Stress and Emotional Availability. Frontiers in psychology 2016;7:1156.

75. Boesveldt S, Graaf K de The Differential Role of Smell and Taste For Eating Behavior. Perception 2017;46:307–319.

76. Lee W, Khan A, Curley JP Major urinary protein levels are associated with social status and context in mouse social hierarchies. Proceedings of the Royal Society B: Biological Sciences 2017;284.

77. Baker M 1,500 scientists lift the lid on reproducibility. Nature 2016;533:452–454.

