## Supplementary figures and images for "Intergenerational effects of a paternal Western diet during adolescence on offspring gut microbiota, stress reactivity and social behavior"

### Supplemental Figure 1

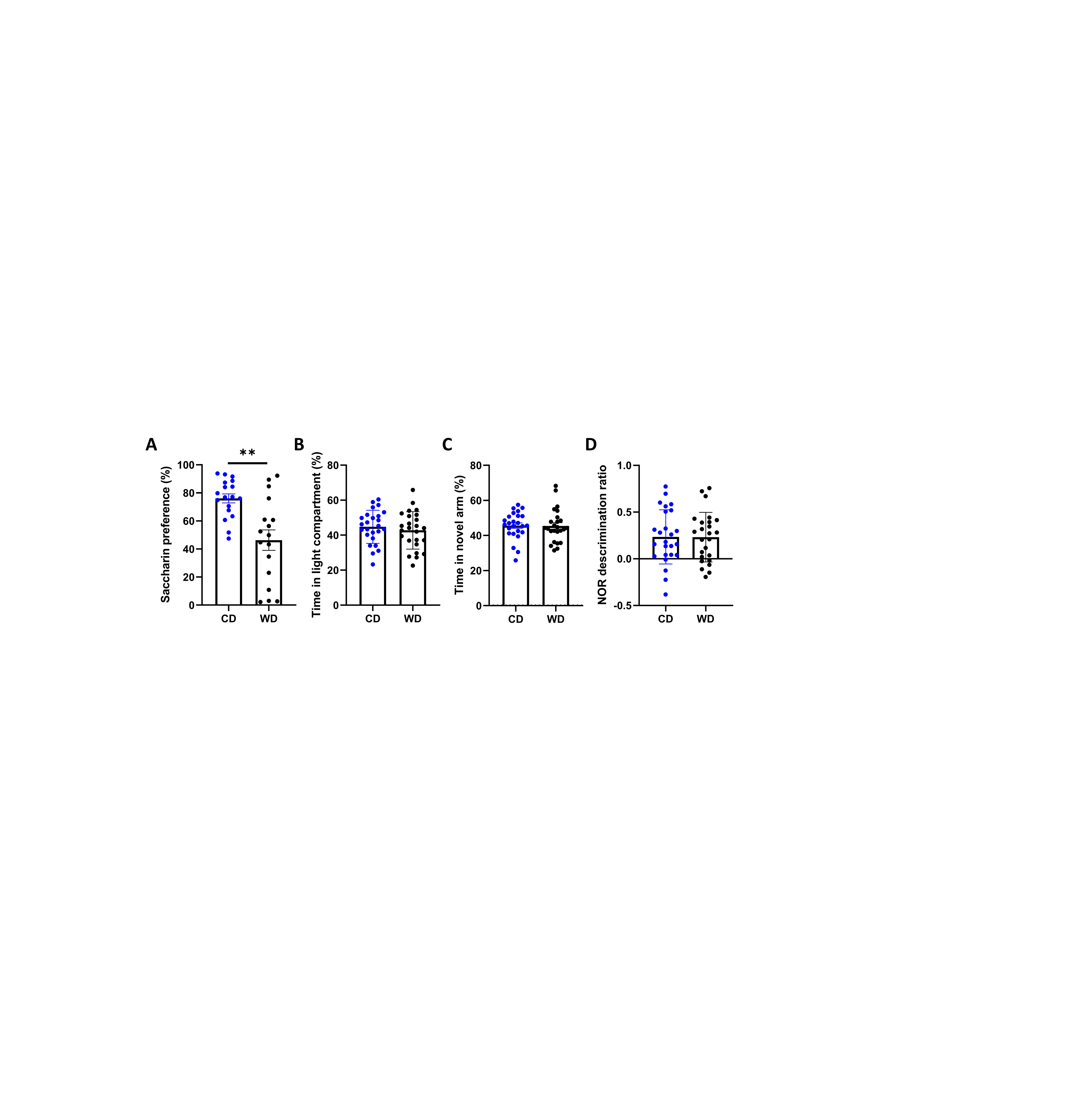

### Supplemental Figure 2

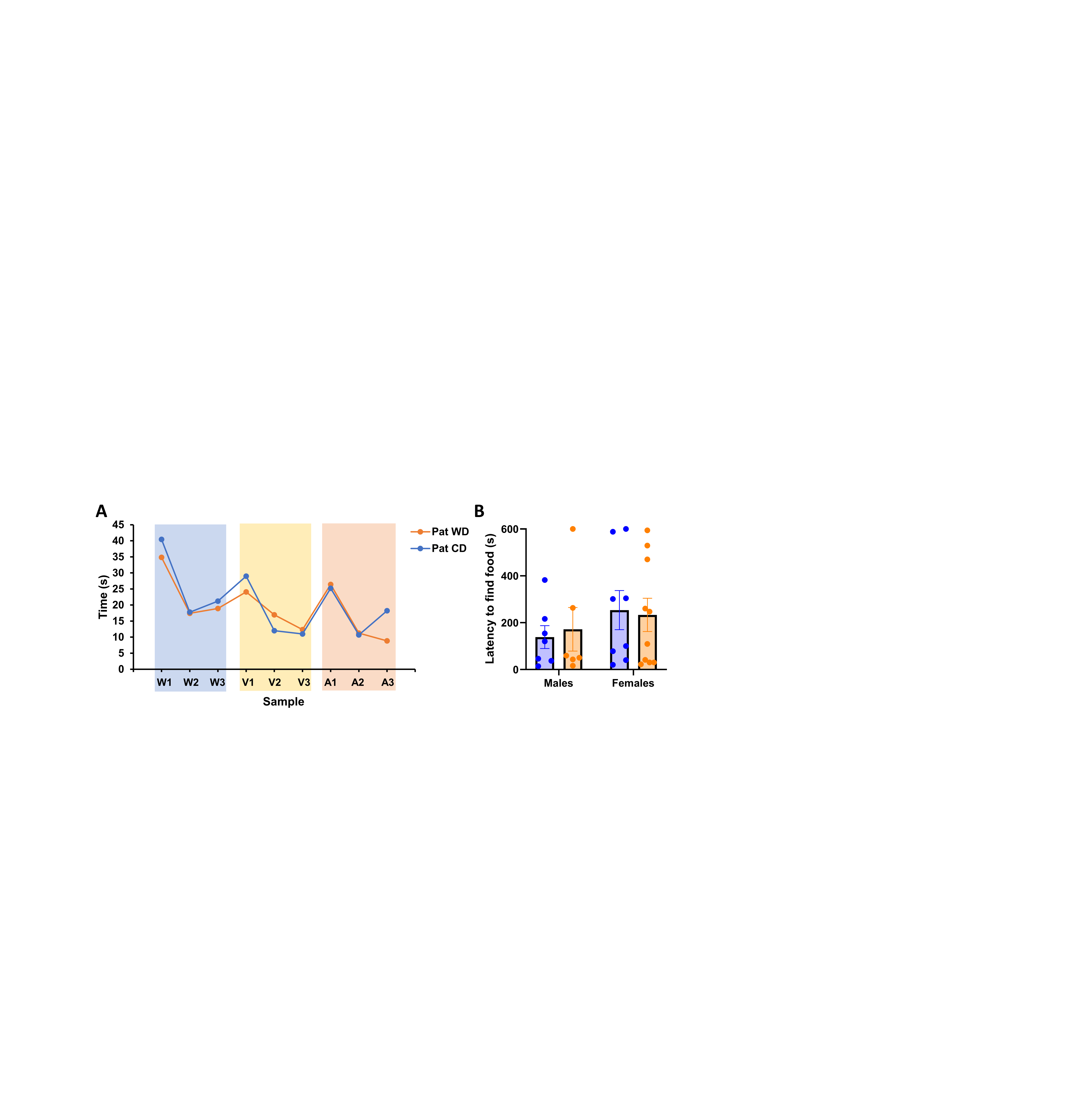
