## Supplemental Table 1 for "Intergenerational effects of a paternal Western diet during adolescence on offspring gut microbiota, stress reactivity and social behavior"

**Supplementary Table 1. Description of maternal behavior patterns.**

| Behavior | Definition |
|---|---|
| Nest building | The mother moves bedding with her snout or her paws while being in a distance to her pups that is smaller than a body length (may occur simultaneously with nursing). |
| Licking and grooming | The mother moves her snout or her paws over the pup, touching it, while her head nods (may occur simultaneously with nursing). |
| Arched-back nursing | The mother is arched over the pups with her legs splayed. |
| Passive nursing | The mother is lying either over the litter but has no arch in her back and there is no obvious extension of her legs, or the mother is lying on her side with one or more pups attached or, more generally, off to the side of the pile of pups. |
| Mother not on nest | The distance between the mother (disregarding the tail) and her pups measures more than a body length. The nest is defined as the area where most of the pups are lying. |
