## Supplemental Table 2 for "Intergenerational effects of a paternal Western diet during adolescence on offspring gut microbiota, stress reactivity and social behavior"

**Supplementary Table 2. Behavioral phenotyping.** Overview of test battery and behavioral aspects and which generation was tested (indicated by an X in the respective column).

| TEST | READOUT | F0 | F1 |
|---|---|---|---|
| Arched-back nursing | High-quality maternal care | X | |
| Buried food test | Olfactory abilities, motivation to find food | | X |
| Elevated plus maze | Anxiety-like behavior, risk-taking behavior | X | X |
| Female preference | Male attractiveness | X | X |
| Licking and grooming | High-quality maternal care | X | |
| Light dark box | Anxiety-like behavior, exploratory locomotion | X | X |
| Novel object recognition | Object memory | X | X |
| Novelty suppressed feeding | Anxiety-like behavior | | X |
| Olfactory discrimination test | Olfactory abilities | | X |
| Passive nursing | Lower-quality maternal care | X | |
| Porsolt swim test | Depression-like behavior | | X |
| Pup retrieval | Maternal care, attentiveness (PND 7) | X | |
| Saccharin preference | Anhedonia | X | X |
| Smell preference | Preference of food based on olfactory cues | | X |
| Social interest and recognition test | Sociability, social memory | | X |
| Surface righting | Motor abilities of pups (PND 7) | | X |
| Taste preference | Preference of food based on taste | | X |
| USV isolation calls | Negative affective state (PND 8) | | X |
| Y-maze | Spatial memory | X | X |
