## Supplemental Table 3 for "Intergenerational effects of a paternal Western diet during adolescence on offspring gut microbiota, stress reactivity and social behavior"

### Supplementary Table 3. Statistical parameters for F0 animals.

| Test - Parameter | Transf. | N | t | df | Sig. (2-tailed) | | Stat. |
|---|---|---|---|---|---|---|---|
| **Elevated plus maze test** | | | | | | | |
| Time on open arms (%) | - | 10 | -0.907 | 18 | 0.376 | | *t* -test |
| Entries into open arms (%) | - | 10 | -0.526 | 18 | 0.605 | | *t* -test |
| Sum of entries (#) | - | 10 | -2.098 | 18 | **0.050** | * | *t* -test |
| **Light-dark test** | | | | | | | |
| Time in light (%) | - | 25 | 0.706 | 48 | 0.484 | | *t* -test |
| Entries into light (#) | Log | 25 | -0.389 | 48 | 0.699 | | *t* -test |
| Latency to enter lit side (s) | Log | 25 | -0.766 | 48 | 0.447 | | *t* -test |
| **Open field test** | | | | | | | |
| Time in centre (s) | - | 10 | -0.329 | 18 | 0.746 | | *t* -test |
| Entries into centre (#) | - | 10 | 0.442 | 18 | 0.664 | | *t* -test |
| Latency to enter centre (s) | Log | 10 | 0.318 | 18 | 0.754 | | *t* -test |
| Total distance (m) | - | 10 | 0.938 | 18 | 0.360 | | *t* -test |
| **Y-maze** | | | | | | | |
| Time in novel arm (%) | - | 25 | 0.077 | 48 | 0.939 | | *t* -test |
| Distance in novel arm (%) | - | 25 | 0.475 | 48 | 0.637 | | *t* -test |
| **Novel object recognition** | | | | | | | |
| Descrimination ratio | - | 25 | 0.035 | 48 | 0.973 | | *t* -test |
| **Female choice test** | | | | | | | |
| w/o bedding: Time in chamber (%) | - | 15 | -1.102 | 28 | 0.280 | | *t* -test |
| w bedding: Time in chamber (%) | - | 20 | 3.019 | 38 | **0.005** | ** | *t* -test |
| w/o mouse: Time in chamber (%) | - | 12 | 4.907 | 22 | **0.000** | *** | *t* -test |
| **Mating behaviour (relative frequencies)** | | | | | | | |
| Approaching | - | 10 | 50.000 | | 1.000 | | *U* -test |
| Following | - | 10 | 44.500 | | 0.676 | | *U* -test |
| Mounting attempts | - | 10 | 47.000 | | 0.819 | | *U* -test |
| Mounting | - | 10 | 11.000 | | **0.003** | ** | *U* -test |
| **Saccharin preference** | | 17-18 | 3.719 | 22.06 | **0.001** | *** | *t* -test |
| **Tissue collections** | | | | | | | |
| Testis weight (mg) | - | 20 | -2.956 | 38 | **0.005** | ** | *t* -test |
| Testis/body weight ratio | - | 20 | 3.045 | 38 | **0.004** | ** | *t* -test |
| Subcutaneous fat (mg) | - | 21 | -10.036 | 33.38 | **0.000** | *** | *t* -test |
| Adipocyte size | - | 5 | 4.983 | 8.000 | **0.000** | *** | *t* -test |
| Epididymal fat (mg) | - | 7-8 | -3.109 | 7 | **0.016** | * | *t* -test |
| Liver (mg) | - | 21 | -4.216 | 40 | **0.000** | *** | *t* -test |
| Liver/body weight ratio | - | 21 | -0.536 | 40 | **0.000** | * | *t* -test |
| Hepatic liver droplets | - | 5 | 2.797 | 5 | **0.042** | * | *t* -test |
| Plasma testosterone | Log | 8 | -0.069 | 14 | 0.946 | | *t* -test |
| Testicular testosterone/g testis | Inv | 8 | -0.304 | 14 | 0.765 | | *t* -test |
| Sperm count | - | 13 | -1.851 | 24 | 0.077 | t | *t* -test |
| **Glucose tolerance test** | | | | | | | |
| t0 | - | 5 | -8.816 | 8 | **0.000** | *** | *t* -test |
| t15 | - | 5 | -11.578 | 8 | **0.000** | *** | *t* -test |
| t30 | - | 5 | -4.536 | 8 | **0.002** | ** | *t* -test |
| t60 | - | 5 | -3.789 | 8 | **0.005** | ** | *t* -test |
| t120 | - | 5 | -1.277 | 8 | 0.237 | | *t* -test |
| **MUPs** | | | | | | | |
| Total MUP concentration | - | 5 | 2.746 | 8 | 0.025 | * | *t* -test |
| MUP/creatinine ratio | - | 5 | 0.756 | 8 | 0.471 | | *t* -test |
| **Body weight** | | | | | | | |
| Food consumption | - | 13 | -6.756 | 24 | **0.000** | *** | *t* -test |
| Energy consumption | - | 13 | -11.49 | 24 | **0.000** | *** | *t* -test |
| PND 28 | - | 25 | 0.908 | 48 | 0.345 | | *t* -test |
| PND 35 | - | 25 | 19.545 | 48 | **0.000** | *** | *t* -test |
| PND 42 | - | 25 | 44.887 | 48 | **0.000** | *** | *t* -test |
| PND 49 | - | 25 | 34.226 | 48 | **0.000** | *** | *t* -test |
| PND 56 | - | 25 | 35.027 | 48 | **0.000** | *** | *t* -test |
| PND 63 | - | 25 | 35.652 | 48 | **0.000** | *** | *t* -test |
| PND 70 | - | 25 | 64.313 | 48 | **0.000** | *** | *t* -test |
| PND 77 | - | 25 | 61.633 | 48 | **0.000** | *** | *t* -test |
| PND 84 | - | 25 | 58.802 | 48 | **0.000** | *** | *t* -test |
| PND 91 | - | 25 | 63.265 | 48 | **0.000** | *** | *t* -test |
| PND 98 | - | 25 | 77.973 | 48 | **0.000** | *** | *t* -test |
| **Pregnancies** | - | 19-21 | 0.129 | 1 | 0.720 | | *t* -test |
| **Maternal care** | | | | | | | |
| Arched-back nursing | - | 10-11 | 2.567 | 19 | **0.019** | * | *t* -test |
| Licking & grooming | - | 10-11 | 2.813 | 19 | **0.011** | * | *t* -test |
| Passive nursing | - | 10-11 | -1.596 | 19 | 0.127 | | *t* -test |
| Active & passive nursing | - | 10-11 | 1.618 | 19 | 0.122 | | *t* -test |
| Not on nest | - | 10-11 | -2.431 | 19 | **0.025** | * | *t* -test |
| **Offspring** | | | | | | | |
| Litter size | - | 16-19 | 1.824 | 33 | 0.077 | t | *t* -test |
| Male offspring | - | 16-19 | 2.342 | 33 | **0.025** | * | *t* -test |
| Female offspring | - | 16-19 | 0.047 | 33 | 0.963 | | *t* -test |
