## Supplemental Table 4 for "Intergenerational effects of a paternal Western diet during adolescence on offspring gut microbiota, stress reactivity and social behavior"

### Supplementary Table 4. Statistical parameters for F1 animals.

| Test - Parameter | Transf. | N | Paternal diet | | | | Sex | | |
|---|---|---|---|---|---|---|---|---|---|
| | | | t/F/X²/U | df | Sig | | t/F/X²/U | df | Sig |
| **USV isolation calls** | | | | | | | | | |
| Latency to call (s) | Inv | 26-31 | 2.868 | 55 | **0.006** | ** | - | - | - |
| Number of calls (#) | Log | 26-31 | 2.278 | 55 | **0.027** | * | - | - | - |
| **Elevated plus maze** | | | | | | | | | |
| Time on open arms (%) | Log | 12-20 | 1.802 | 64 | 0.184 | | 0.390 | 64 | 0.534 |
| Entries into open arms (%) | Log | 12-20 | 3.224 | 64 | 0.077 | | 0.877 | 64 | 0.353 |
| Sum of entries (#) | Log | 12-20 | 0.362 | 64 | 0.549 | | 0.967 | 64 | 0.329 |
| **Light-dark box** | | | | | | | | | |
| Time in light (%) | - | 22-32 | 0.951 | 111 | 0.332 | | 0.012 | 111 | 0.914 |
| Latency to enter light (s) | - | 22-32 | 0.003 | 111 | 0.957 | | 8.006 | 111 | **0.006** |
| Entries into light (#) | SqRt | 22-32 | 0.446 | 111 | 0.506 | | 3.922 | 111 | 0.050 |
| Total distance (m) | - | 22-32 | 0.183 | 111 | 0.670 | | 6.086 | 111 | **0.015** |
| **Y-maze** | | | | | | | | | |
| Time in novel arm (%) | - | 9-16 | 3.654 | 47 | 0.062 | | 1.209 | 47 | 0.277 |
| Distance in novel arm (%) | - | 9-16 | 4.225 | 47 | **0.045** | * | 5.861 | 47 | **0.019** |
| **Novel object recognition** | | | | | | | | | |
| Recognition index | - | 9-16 | 1.439 | 47 | 0.236 | | 0.763 | 47 | 0.387 |
| **Social interaction and recognition** | | | | | | | | | |
| Time in social interaction zone (%) | - | 8-11 | 0.066 | 34 | 0.799 | | 0.515 | 34 | 0.478 |
| Time in novel interaction zone (%) | - | 8-11 | 0.504 | 34 | 0.111 | | 0.369 | 34 | 0.548 |
| **Novelty-suppressed feeding** | | | | | | | | | |
| Latency to start eating (s) | - | 12-16 | 0.158 | 50 | 0.692 | | 3.088 | 50 | 0.085 |
| Food choice | - | 12-16 | 12.208 | 1 | **0.000** | *** | 0.021 | 1 | 0.884 |
| **Food smell preference** | | | | | | | | | |
| Time sniffing sample (s) | - | 8-11 | 0.128 | 37 | 0.723 | | 0.054 | 37 | 0.818 |
| **Food taste preference (48 h)** | | | | | | | | | |
| WD preference (%) | * | 7-10 | 21.000 | | 0.172 | | 23.000 | | 0.242 |
| **Olfactory discrimination** | | | | | | | | | |
| Time sniffing sample (s) | - | 9-12 | 0.209 | 15 | 0.654 | | 0.376 | 1 | 0.549 |
| **Buried food test** | | | | | | | | | |
| Latency to find food (s) | Log | 6-10 | 0.050 | 27 | 0.824 | | 0.989 | 27 | 0.329 |
| **Porsolt swim test** | | | | | | | | | |
| Latency to immobility (s) | Log | 13-17 | 23.215 | 55 | **0.000** | *** | 0.180 | 55 | 0.673 |
| Time immobile (s) | - | 7-8 | 0.861 | 26 | 0.362 | | 0.494 | 26 | 0.488 |
| **Dominance tube test** | - | 10-11 | -5.134 | 19 | **0.000** | *** | - | - | - |
| **Saccharin preference** | - | 15-21 | 0.021 | 69 | 0.886 | | 0.063 | 69 | 0.802 |
| **Female choice test** | | | | | | | | | |
| w bedding: Time in chamber (%) | - | 14-16 | -5.563 | 28 | **0.000** | *** | - | - | - |
| w/o mouse: Time in chamber (%) | - | 14-16 | -5.597 | 28 | **0.000** | *** | - | - | - |
| **Tissue collections** | | | | | | | | | |
| Liver (mg) | - | 7-12 | 1.707 | 33 | 0.200 | | 31.556 | 33 | **0.000** |
| Liver/body weight ratio | - | 7-12 | 0.785 | 33 | 0.382 | | 0.321 | 33 | 0.575 |
| Subcutaneous fat (mg) | | 9-12 | -1.415 | 19 | 0.173 | | - | - | - |
| Testis weight (mg) | - | 12-15 | -0.375 | 25 | 0.711 | | - | - | - |
| Testis/body weight ratio | - | 12-15 | 0.509 | 25 | 0.615 | | - | - | - |
| Sperm count | - | 11-14 | 1.461 | 23 | 0.157 | | - | - | - |
| **Endocrinological parameters** | | | | | | | | | |
| Plasma testosterone | Inv | 15-16 | -3.129 | 17.04 | **0.006** | ** | - | - | - |
| Testicular testosterone/g testis | Log | 7-11 | 0.624 | 16 | 0.535 | | - | - | - |
| Corticosterone (baseline) | - | 6-10 | 0.350 | 28 | 0.559 | | 6.413 | 28 | **0.017** |
| Corticosterone (reaction) | - | 9-12 | 0.585 | 37 | 0.449 | | 2.133 | 37 | 0.153 |
| **Body weight (g)** | | | | | | | | | |
| PND 7 | - | 39-56 | 6.943 | 201 | **0.009** | ** | 2.732 | 201 | 0.100 |
| PND 14 | - | 39-56 | 23.257 | 201 | **0.000** | *** | 0.279 | 201 | 0.598 |
| PND 21 | - | 39-56 | 20.924 | 201 | **0.000** | *** | 9.488 | 201 | **0.002** |
| PND 28 | | 39-56 | 0.801 | 201 | 0.372 | | 52.528 | 201 | **0.000** |
| PND 35 | - | 39-56 | 0.674 | 201 | 0.413 | | 232.716 | 201 | **0.000** |
| PND 42 | - | 28-40 | 0.027 | 143 | 0.869 | | 335.851 | 143 | **0.000** |
| PND 49 | - | 21-34 | 0.009 | 110 | 0.923 | | 385.502 | 110 | **0.000** |
| PND 56 | - | 21-34 | 0.022 | 110 | 0.883 | | 476.645 | 110 | **0.000** |
| PND 63 | - | 21-34 | 1.751 | 110 | 0.188 | | 430.605 | 110 | **0.000** |
| PND 70 | - | 15-26 | 0.533 | 110 | 0.468 | | 392.151 | 110 | **0.000** |
| PND 77 | - | 11-22 | 0.011 | 63 | 0.916 | | 455.910 | 63 | **0.000** |
| PND 84 | - | 15-26 | 0.063 | 79 | 0.802 | | 468.731 | 79 | **0.000** |
| PND 91 | - | 15-26 | 0.136 | 79 | 0.714 | | 493.314 | 79 | **0.000** |
| PND 98 | - | 11-21 | 0.187 | 62 | 0.667 | | 383.557 | 62 | **0.000** |
| PND 105 | - | 11-22 | 0.73 | 63 | 0.396 | | 417.286 | 63 | **0.000** |
| **16S alpha diversity** | | | | | | | | | |
| Observed | Inv | 12 | 0.001 | 20 | 0.976 | | - | - | - |
| Shannon | - | 12 | 0.727 | 20 | 0.404 | | - | - | - |
| Inverse Simpson | - | 12 | 1.326 | 20 | 0.263 | | - | - | - |
| **16S beta diversity** | | | | | | | | | |
| R2 | - | 12 | 0.033 | 20 | 0.673 | | - | - | - |
| **16 S differential abundance with ANCOM** | | | | | | | | | |
| Actinobacteria | Log | 12 | 9.868 | 20 | **0.005** | ** | - | - | - |
| **Body composition** | | | | | | | | | |
| Fat | - | 6-10 | 0.004 | 28 | 0.952 | | 23.672 | 28 | **0.000** |
| Lean | - | 6-10 | 0.042 | 28 | 0.838 | | 82.566 | 28 | **0.000** |
| Fluid | - | 6-10 | 1.662 | 28 | 0.208 | | 8.798 | 28 | **0.006** |
| **Metabolic cages** | | | | | | | | | |
| Respiratory quotient | - | 6-10 | 1.742 | 28 | 0.198 | | 18.223 | 28 | **0.000** |
| Energy expaditure | - | 6-10 | 0.740 | 28 | 0.397 | | 20.490 | 28 | **0.000** |
| Water vapor loss | - | 6-10 | 0.131 | 28 | 0.720 | | 2.662 | 28 | 0.114 |
| **MUPs** | | | | | | | | | |
| Total MUP concentration | - | 6-9 | 1.123 | 13 | 0.282 | | - | - | - |
| MUP/creatinine ratio | - | 6-9 | -0.556 | 13 | 0.587 | | - | - | - |
